# CryoEM structure of an MHC-I/TAPBPR peptide bound intermediate reveals the mechanism of antigen proofreading

**DOI:** 10.1101/2024.08.05.606663

**Authors:** Yi Sun, Ruth Anne Pumroy, Leena Mallik, Apala Chaudhuri, Chloe Wang, Daniel Hwang, Julia N. Danon, Kimia Dasteh Goli, Vera Moiseenkova-Bell, Nikolaos G. Sgourakis

## Abstract

Class I major histocompatibility complex (MHC-I) proteins play a pivotal role in adaptive immunity by displaying epitopic peptides to CD8+ T cells. The chaperones tapasin and TAPBPR promote the selection of immunogenic antigens from a large pool of intracellular peptides. Interactions of chaperoned MHC-I molecules with incoming peptides are transient in nature, and as a result, the precise antigen proofreading mechanism remains elusive. Here, we leverage a high-fidelity TAPBPR variant and conformationally stabilized MHC-I, to determine the solution structure of the human antigen editing complex bound to a peptide decoy by cryogenic electron microscopy (cryo-EM) at an average resolution of 3.0 Å. Antigen proofreading is mediated by transient interactions formed between the nascent peptide binding groove with the P2/P3 peptide anchors, where conserved MHC-I residues stabilize incoming peptides through backbone-focused contacts. Finally, using our high-fidelity chaperone, we demonstrate robust peptide exchange on the cell surface across multiple clinically relevant human MHC-I allomorphs. Our work has important ramifications for understanding the selection of immunogenic epitopes for T cell screening and vaccine design applications.

## Introduction

Class I major histocompatibility complex (MHC-I) proteins display the intracellular proteome onto the cell surface for immunosurveillance by sampling a large pool of peptide fragments^1–3^. Optimal high-affinity-peptide-loaded MHC-I (pMHC-I) molecules are recognized by T cell receptors (TCR) and natural killer (NK) receptors, triggering downstream cellular activation and clonal expansion and resulting in the clearance of infected, or aberrant cells^4–6^. While the immunogenic peptide repertoires of the Class I Human Leukocyte Antigen (HLA, human MHC-I) proteins are defined by their extremely polymorphic grooves^7–9^, selection and optimization of the repertoires in the cellular pathway are mediated by two dedicated molecular chaperones, tapasin^10–12^ and TAP binding protein-related (TAPBPR)^13,14^. TAPBPR is a homolog of tapasin that is not part of the peptide loading complex (PLC). These chaperones serve as an essential quality control checkpoint for pMHC-I molecules, optimizing the peptide cargo of the MHC-I molecules *en route* to the cell surface^15,16^. Polymorphic residues at the MHC-I groove and chaperone interaction surfaces confer a wide range of dependence on chaperones for peptide loading and cell surface expression across various HLA allotypes, with important ramifications for viral control among different individuals^17–21^. In addition, these antigen processing and presentation (APP) chaperones have recently emerged as key components for the presentation of metabolite ligands on MHC-related 1 (MR1) molecules^22–24^, and have provided powerful tools for *in vitro* applications, including the generation of pMHC-I libraries encompassing different peptide specificities or the loading of antigens on cells independently of the endogenous processing pathway^18,25–27^.

Two independently solved crystal structures of mouse MHC-I bound to human TAPBPR^28,29^, together with studies of TAPBPR and tapasin in complex with human MHC-I molecules by X-ray crystallography, nuclear magnetic resonance (NMR) spectroscopy, and cryogenic electron microscopy (cryo-EM)^30–32,21,33–35^ have provided key insights into the chaperoning mechanism. Empty MHC-I adopts an open, peptide-receptive conformation where the short α_2-1_ helix shifted outwards by 3 Å to induce a widened peptide binding groove^28,29^. Based on these structures obtained for the empty, resting state of the complex, it has been proposed that the binding of high-affinity peptides triggers the closure of the MHC-I groove, which allosterically promotes the release of pMHC-I from the chaperone^28,29,32^. A range of biophysical studies have also demonstrated that the affinity of TAPBPR for MHC-I is negatively correlated with peptide occupancy^18,21,31^. TAPBPR preferentially interacts with nascent empty or suboptimal peptide-loaded MHC-I molecules, exchanging low- for high-affinity peptides. Therefore, the precise molecular mechanism of MHC-I antigen selection by TAPBPR remains enigmatic^36,37^ due to the transient nature of the peptide/MHC-I/TAPBPR complex, resulting in no structural visualization and precluding a detailed understanding of how TAPBPR edits the peptide repertoire.

To examine the editing function of TAPBPR and capture the biologically relevant peptide-bound state of its MHC-I complex, we have leveraged protein engineering to enhance the stability of MHC-I molecules that are loaded with suboptimal low-to-moderate-affinity peptides^38^, as well as deep scanning mutagenesis of MHC-I binding surfaces on TAPBPR^18^. Here, we characterize a high-fidelity chaperone, named TAPBPR^HiFi^, which shows enhanced binding to peptide-loaded MHC-I molecules and peptide exchange function *in vitro*. We determine the cryoEM structure of TAPBPR^HiFi^/HLA-A*02:01/β_2_m loaded with a peptide decoy, revealing a key intermediate of the peptide exchange process. We find that the transient proofreading complex utilizes (1) conserved groove residues to capture the backbone of incoming peptides in a native-like conformation, (2) properly conformed hydrophobic A-, B-, and D-pockets to dock the peptide P2 and P3 anchors, and (3) a conformationally heterogenous F-pocket that folds upon binding to high-affinity peptides, allosterically triggering the dissociation of TAPBPR. Finally, we show that TAPBPR^HiFi^ significantly enhances peptide exchange across multiple HLA-A allotypes expressed on a cellular membrane to promote antigen loading independently of the endogenous pathway, which involves tapasin and the peptide-loading complex, with important applications in screening for and eliciting T cell responses in various experimental and therapeutic settings. Taken together, our combined structural, biochemical, and functional studies provide a complete mechanism of MHC-I antigen proofreading and peptide repertoire selection.

## Results

### Enhanced editing of MHC-I antigens using an engineered molecular chaperone

While contradictory peptide unloading/levering^39,40^ or trapping^41^ mechanisms have been proposed involving the G24-R36 loop of TAPBPR, whether the loop actively participates in cargo editing of the peptide-loaded MHC-I molecules remains unclear^36^. We sought to resolve this controversy and identify TAPBPR regions that promote interactions with peptide-loaded molecules for antigen editing. Towards this, we leveraged either the wildtype TAPBPR, TAPBPR^WT^, or our engineered version high-fidelity TAPBPR^HiFi^, containing 3 mutations S104F, K211L, and R270Q outside the loop region^18^, to directly measure binding to a high-affinity peptide TAX9 loaded HLA-A*02:01 by surface plasmon resonance (SPR). Experiments were run in the presence of TAX9 peptide in large molar excess to prevent peptide dissociation upon binding to TAPBPR. We also compared different mutants of TAPBPR^HiFi^ and TAPBPR^WT^, containing partial (A29-S32 deletion, TAPBPR^ΔALAS^) and complete loop deletions (G24-R36 deletion, TAPBPR^ΔG24-R36^ and TAPBPR^HiFiΔG24-R36^) as well as TAPBPR^TN6^ (E205K, R207E, Q209S, and Q272S), which does not interact with peptide-loaded MHC-I (**Figure 1A and S1**)^42,31,26^. Our results showed that TAX9-loaded HLA-A*02:01 exhibited an enhanced affinity (reduced K_D_ by one order of magnitude) for TAPBPR^HiFi^, relative to TAPBPR^WT^ (**Figure 1B-C and S2**). Meanwhile, loop deletions exhibited no significant effect for TAPBPR^HiFi^ but an approximately 3-fold decrease in K_D_ for TAPBPR^WT^ on binding to pMHC-I (**Figure 1B-C and S2**), likely due to the loss of hydrophobic interactions with the rim of the MHC-I α_1_ and α_2_ helices, in agreement with our previous mapping studies^39,41^. These observations demonstrate that TAPBPR^HiFi^ improves the recognition of peptide-loaded MHC-I molecules and tolerates modifications, compared to the wild type.

**Figure 1.**
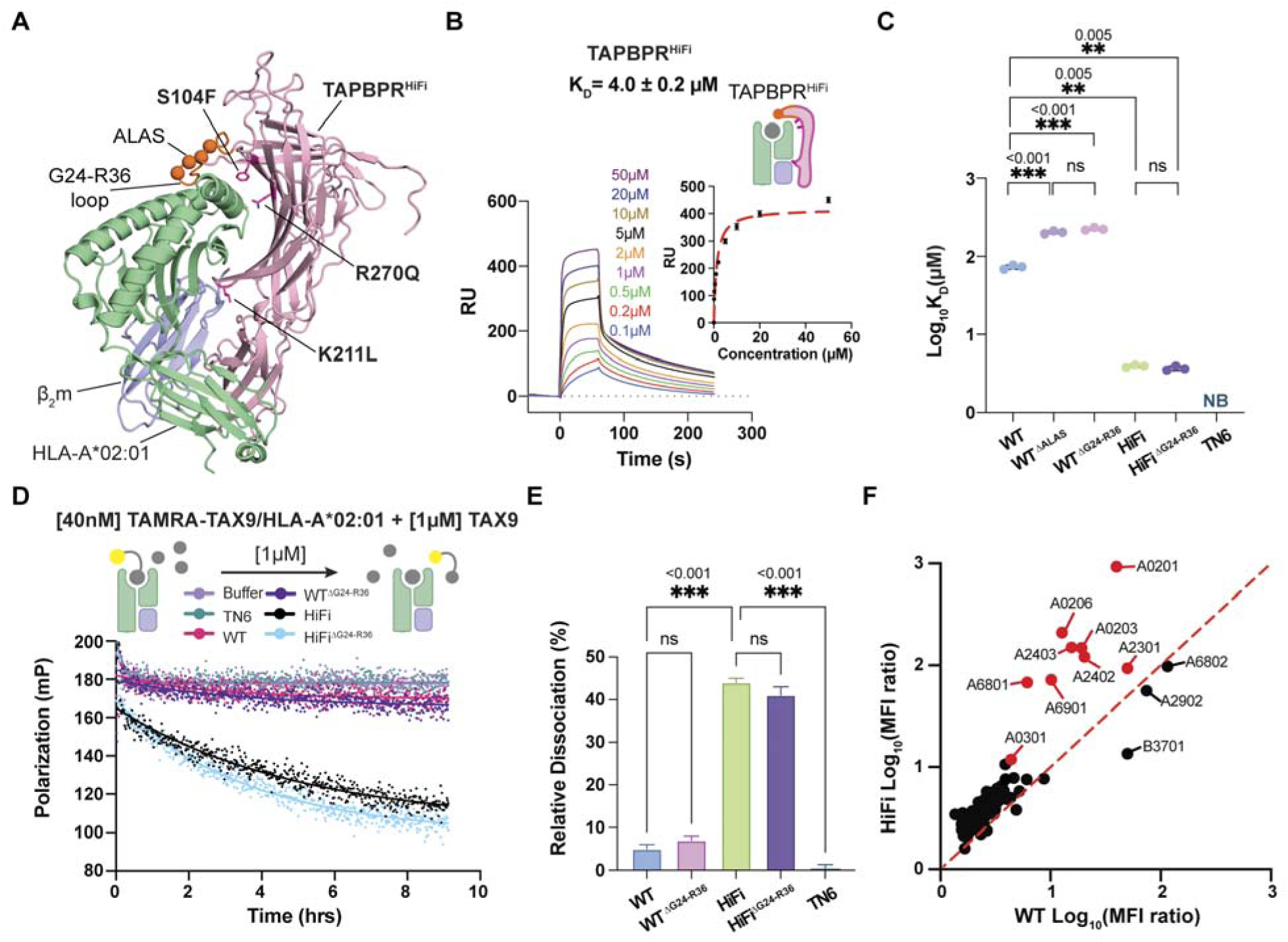
A high-fidelity TAPBPR variant TAPBPR^HiFi^ enhances peptide editing on peptide-loaded MHC-I. (**A**) Structural model of TAPBPR^HiFi^ in complex with peptide-free HLA-A*02:01/β2m generated using RosettaCM^56^. The G24-R36 loop is colored orange, and the A29-S32 loop segment is shown as spheres. The sidechains of mutations S104F, K211L, and R270Q are shown as magenta sticks. (**B**) Representative SPR sensorgram of graded concentrations of TAPBPR^HiFi^ flown over a streptavidin chip coupled with TAX9/HLA-A*02:01 in excess TAX9 peptide. (**C**) Log-scale comparison of K_D_ values between TAX9/HLA-A*02:01 and TAPBPR. Results of three independent experiments (mean ± SD) are shown as scatter plots. (**D**) Peptide dissociation kinetics of fluorophore-labeled TAMRA-TAX9-(TAMRA-KLFGYPVYV)-peptide-loaded HLA-A*02:01 in the presence of excess unlabeled TAX9 peptide with buffer (light purple), TAPBPR^TN6^ (green), TAPBPR^WT^ (magenta), TAPBPR^ΔG24-R36^ (purple), TAPBPR^HiFi^ (light blue) and TAPBPR^HiFiΔG24-R36^ (black). Data are means for n = 3 independent experiments. (**E**) Relative peptide dissociation of TAMRA-TAX9 from HLA-A*02:01 by TAPBPR relative to no TAPBPR. The relative dissociation was calculated using the equation % 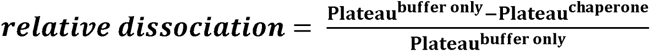, and the plateau for each dissociation was individually extracted by fitting one phase decay. Error bars (SD) were propagated from three independent experiments. (**F**) Comparison of binding profiles for TAPBPR^WT^ and TAPBPR^HiFi^ against a panel of 97 common HLA allotypes using single antigen beads (SABs). Two-sample unequal variance student’s t-test was performed, P > 0.12 (not significant, ns), P < 0.033(*), P < 0.002(**), and P < 0.001(***). See also **Figure S1-S3**.

We hypothesized that enhanced TAPBPR binding to peptide-loaded MHC-I molecules should correlate with a promotion of peptide dissociation *in vitro*. We then applied a fluorescence polarization (FP) assay employing HLA-A*02:01 refolded with a TAMRA-labeled TAX9 peptide to directly assess peptide unloading from the MHC-I^38^. We found that TAPBPR^WT^ and its loop deletion mutant, TAPBPR^ΔG24-R36^, demonstrated a similarly low level of peptide unloading, relative to the negative controls (**Figure 1D-E**). Notably, both TAPBPR^HiFi^ and TAPBPR^HiFiΔG24-R36^ significantly improved peptide unloading function (**Figure 1D-E**). However, introducing G24-R36 loop deletion on either TAPBPR^WT^ or TAPBPR^HiFi^ had no functional impact on MHC-I peptide editing *in vitro* (**Figure 1E**). Taken together, our results suggest that, while the G24-R36 loop has an important role in maintaining TAPBPR’s structural integrity for pMHC-I recognition, surfaces outside the loop are essential for promoting interactions with peptide-loaded MHC-I and enhancing the editing capability of TAPBPR^18^.

To determine the MHC-I allelic interaction landscape of TAPBPR^HiFi^, we applied a single antigen bead (SAB) assay encompassing 97 common HLA allotypes^18,43^. We measured the phycoerythrin (PE) mean fluorescent intensity (MFI) of SABs upon incubation with TAPBPR^HiFi^ or TAPBPR^WT^ PE-tetramers. The levels of nonspecific background binding to HLAs are corrected by the corresponding TAPBPR PE-tetramer staining of SABs that are pre-incubated with W6/32, a pan-allelic HLA monoclonal antibody, which blocks the TAPBPR interaction surface on MHC-I, and further compared to the staining of TAPBPR^TN6^ PE-tetramer (**Figure S3A-B**). Analysis of MFI ratios for TAPBPR^HiFi^ revealed enhanced interactions with multiple HLA allotypes, including HLA-A*02:01, A*02:03, A*02:06, A*69:01, A*68:01, A*23:01, A*24:02, and A*24:03, relative to TAPBPR^WT^ (**Figure 1F and S3C**). Together, these results demonstrate that a broad scope of HLA-A allotypes are prone to enhanced recognition by TAPBPR^HiFi^, which can further promote peptide editing, indicating its key role in the cellular pathway for MHC-I antigen selection^26^.

### Solution mapping of TAPBPR^HiFi^ interactions with peptide-loaded MHC-I

We, therefore, ask how TAPBPR^HiFi^ acts on properly conformed, peptide-loaded MHC-I compared to TAPBPR^WT^ in a solution environment to achieve enhanced binding affinity and editing function. We used 2D NMR spectroscopy to elucidate the modes of binding and dynamics of the TAPBPR^HiFi^ interactions with high-affinity peptide TAX9-loaded HLA-A*02:01. By titrating unlabeled TAPBPR^HiFi^ on isotopically methyl-labeled (AILV) HLA-A*02:01 using established methods^21,32,41^, 42 out of 96 resonances undergo changes in the slow exchange regime, which is similar to the changes upon binding to TAPBPR^WT^ (**Figure 2A and S4A**). We observed a complete shift in the population to a chaperone-bound form for nearly all resonances at a three-fold molar excess of TAPBPR^HiFi^ (**Figure 2B and S4A-B**), indicating the presence of a high-affinity complex. Analysis of methyl ^13^C/^1^H chemical shift deviations (CSDs) captures the residues that experience significant conformational changes (≥0.5SD) based on the difference in the local magnetic environment (**Figure 2C**). Comparing the CSDs of TAX9-loaded HLA-A*02:01 when binding to TAPBPR^HiFi^ relative to TAPBPR^WT^ results in a Pearson correlation coefficient of 0.93 (**Figure 2D**), suggesting that TAX9/HLA*02:01 shifts to a similar conformational state when bound to TAPBPR^HiFi^ or TAPBPR^WT^. Taken together, TAPBPR^HiFi^ docks on MHC-I using an overall similar binding mode, in comparison to our previous solution mapping of interactions between TAX9/HLA-A*02:01 with the wildtype TAPBPR^21,41^.

**Figure 2.**
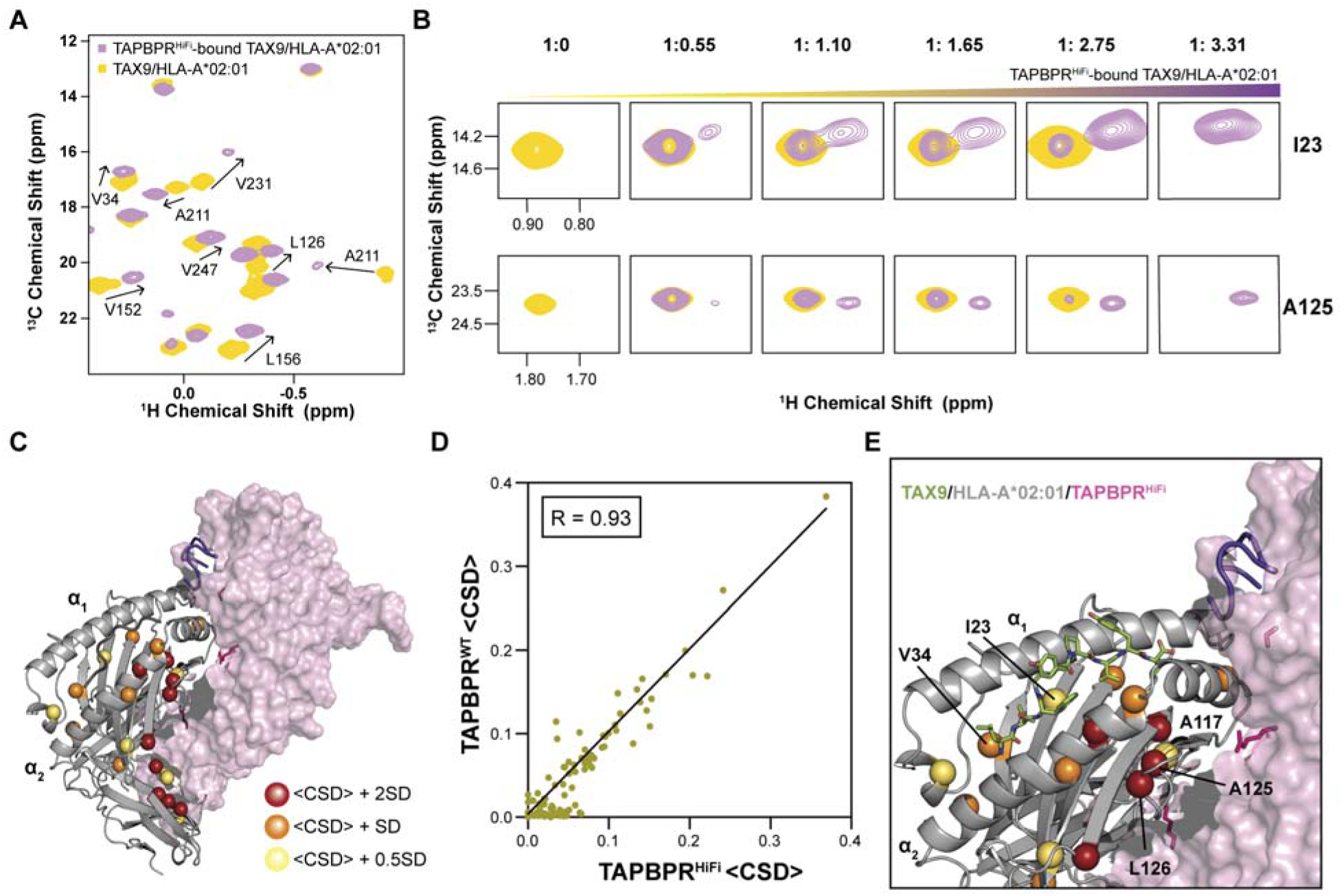
TAPBPR^HiFi^ uses a conserved docking mode to recognize peptide-loaded HLA-A*02:01 in solution. (**A**) A representative region of the 2D ^1^H-^13^C HMQC spectral overlay of selectively ^13^C/^1^H AILV methyl-labeled HLA-A*02:01/TAX9/β_2_m (at the heavy chain) on a ^12^C/^2^H background (yellow) and with a 3-fold molar excess of TAPBPR^HiFi^ at natural isotopic abundance (purple). (**B**) Selected NMR resonances (corresponding to the methyl sidechains of I23 and A125) undergoing conformational exchange between the free and TAPBPR^HiFi^-bound states. The HLA-A*02:01: TAPBPR^HiFi^ molar ratios for the different titration points were 1: 0, 1: 0.55, 1: 1.10, 1: 1.65, 1: 2.75 and 1: 3.31. (**C**) Methyl groups of residues undergoing significant chemical shift deviation (CSD) upon binding of TAPBPR^HiFi^ are mapped onto the structure of the HLA-A*02:01 (gray)/TAPBPR^HiFi^ (pink) complex. The structure is a RosettaCM^56^ homology model obtained using the H2-D^d^/TAPBPR crystal structure as a template (PDB ID: 5WER). CSDs are plotted using a heat-map scale shown on the right. (**D**) Correlation plot of CSDs observed for the titration of HLA-A*02:01 with TAPBPR^WT^^21,41^ versus TAPBPR^HiFi^. The Pearson correlation coefficient (r) is shown on the plot (P value < 0.0001). (**E**) Close-up of the peptide-binding groove with the TAX9 peptide (shown as green sticks). Select residues distributed throughout the MHC-I groove and their methyl CSDs are labeled. The three mutation sites on TAPBPR^HiFi^ are denoted as pink spheres. See also **Figure S4**.

Our NMR data showed that residues, such as A117, A125, and L126, located near the TAPBPR contacting surface demonstrated significant CSDs (**Figure 2F**). Multiple residues like I23 and V34 that are located more than 10 Å away from the interface or within the peptide binding groove also exhibited significant CSDs (**Figure 2F**), indicating allosteric, long-range effects on pMHC-I by TAPBPR^HiFi^. Next, we used a line shape analysis of our 2D methyl HSQC spectra to determine the affinity between TAPBPR^HiFi^ and high-affinity peptide-loaded HLA-A*02:01 without immobilization. Consistent with our previous SPR measurements, TAPBPR^HiFi^ demonstrated a low micromolar-range affinity (K_D_ = 13.9 μM) (**Table S1)** when fitting the resonances that undergo conformational changes to a two-state binding model (**Figure S4C**). Together, these observations indicate that pMHC-I molecules experience similar structural changes upon binding to TAPBPR^HiFi^ or TAPBPR^WT^ and yet form a high-affinity complex with TAPBPR^HiFi^. This motivates us to use TAPBPR^HiFi^ as a bait to stabilize MHC-I in a peptide editing state and capture the MHC-I/TAPBPR proofreading complex.

### CryoEM structure of the MHC-I/TAPBPR complex bound to a peptide decoy

Next, we reasoned that the enhanced affinity of TAPBPR^HiFi^ for pMHC-I could capture transient, unstable peptide-editing complexes that would not be resolvable with TAPBPR^WT^. To define the structural basis of peptide antigen proofreading and selection by TAPBPR, we leveraged our enhanced peptide editor, TAPBPR^HiFi^, to isolate a tertiary complex prepared with open HLA-A*02:01 (G120C) refolded with β_2_m (H31C)^38^ and a photocleavable peptide (KILGFVFJV, J = 3-amino-3-(2-nitrophenyl)-propionic acid) used as a conditional ligand^44,45^. Upon UV irradiation, we purified soluble TAPBPR^HiFi^ in complexes with open HLA-A*02:01/β_2_m by size exclusion chromatography (SEC) (**Figure 3A**). Further analysis by LC-MS revealed that a 7-mer peptide decoy (KILGFVF) was captured within the TAPBPR/MHC-I complex (**Figure 3B and S5**). We solved the structure of this purified 90 kDa complex using cryoEM (**Table S2**). Briefly, purified peptide-loaded open HLA-A*02:01/TAPBPR^HiFi^ was used to screen grids, and more than 5,000 micrographs were collected (**Figure S6**). This dataset revealed a fully assembled MHC-I/TAPBPR complex with a well-defined peptide binding groove, though with some heterogeneity in peptide occupancy. We sorted the dataset to obtain a set of particles with the best occupancy of the first residues of our peptide decoy, obtaining a structure at an overall resolution of 3.0 Å (**Figure S7-8**).

**Figure 3.**
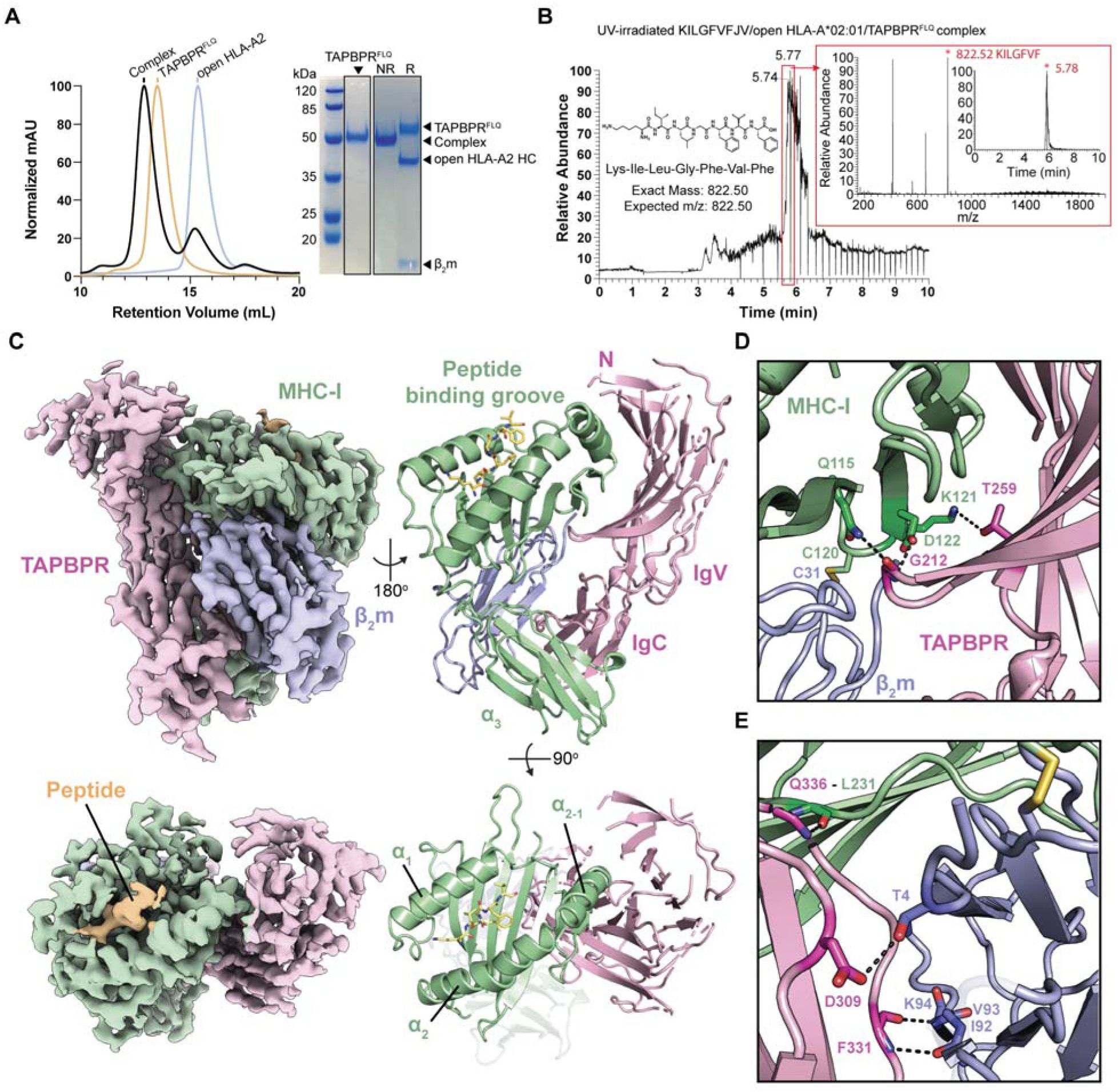
Isolation and cryoEM structure of an MHC-I/TAPBPR complex bound to a peptide decoy. (**A**) SEC purification of a recombinant open HLA-A*02:01/TAPBPR^HiFi^ complex prepared by UV-irradiation of KILGFVFJV/HLA-A*02:01 pre-incubated with TAPBPR^HiFi^ at 1:1.3 molar ratio. J = 3-amino-3-(2-nitrophenyl)-propionic acid. Sodium Dodecyl Sulphate/Polyacrylamide Gel Electrophoresis analysis confirms the identity of the TAPBPR^HiFi^/open HLA-A*02:01 complex peak under nonreducing (NR) or reducing (R) conditions. The bands of TAPBPR^HiFi^, open HLA-A*02:01 heavy chain (HC), and β_2_m, as well as the positions of molecular weight standards, are indicated. (**B**) LC/MS analysis of the purified complex in Fig. 3A showing the presence of captured peptide decoy KILGFVF (observed and expected mass-to-charge ratios are 822.52 and 822.50 m/z, respectively). (**C**) Reconstructed cryoEM density and refined structural model of the tertiary complex, with TAPBPR^HiFi^ shown in light pink, open HLA-A*02:01 heavy chain in pale green, β_2_m in light blue, and peptide in wheat, as indicated. (**D**) Zoom in on the interactions between the TAPBPR^HiFi^ hairpin and the floor of the MHC-I peptide binding groove. (**E**) Zoom in on the interactions among TAPBPR^HiFi^ IgC domain residues, MHC-I α_3_, and β_2_m, highlighting the sidechain interaction between TAPBPR^HiFi^ and β_2_m. See also **Figure S5-S8**.

Inspection of our structural model built into well-defined regions of the electron density reveals key chaperoning interactions between peptide-loaded MHC-I and TAPBPR (**Figure 3C**). In agreement with previous studies^28,29^, we observe that the N-terminal immunoglobulin V (IgV)– like domain of TAPBPR^HiFi^ cradles the pMHC-I peptide binding groove, while the C-terminal IgC domain nestles between the pMHC-I α_3_and β_2_m domains to create a pseudo 3-fold symmetric arrangement (**Figure 3C**). The overall binding mode of TAPBPR is polarized towards the α_2_ helix of the pMHC-I peptide binding groove (**Figure 3C**), consistent with the crystal structures of empty mouse MHC-I/TAPBPR complexes^28,29^ as well as extensive mapping of human MHC-I/TAPBPR interactions by NMR, hydrogen-deuterium exchange, and deep scanning mutagenesis^18,21,32^. Interactions between TAPBPR and the groove of pMHC-I are mediated through G212 and T259, forming polar contacts with Q115, K121, and D122 on strands β7 and β8, located under the peptide binding site (**Figure 3D**). We also notice interactions between TAPBPR residues Q336 and F331 with L231 from α_3_ and residues I92-K94 from β_2_m, while D309 of TAPBPR forms a hydrogen bond with the T4 sidechain from β_2_m (**Figure 3E**). In summary, a triad of domain interactions between β_2_m, pMHC-I α_3,_ and C terminal TAPBPR mediate stable docking, and allow TAPBPR to stabilize the floor and α_2-1_ helix of the peptide binding groove for the peptide-loaded MHC-I molecules, setting the stage for peptide editing.

### Peptide-loaded MHC-I/TAPBPR complex reveals unexpected structural alterations

To understand the molecular mechanism of peptide editing and conformational changes induced by peptide binding to the MHC-I/TAPBPR proofreading complex, we performed a structural comparison of our partially loaded KILGFVF/HLA-A*02:01/TAPBPR cryoEM structure relative to the fully loaded HLA-A*02:01/GILGFVFTL X-ray structure (**Figure 4A, Movie S1**)^46^. We find that the α_2-1_ helix, comprising residues E148-H151, adopts a conformation that is widened by approximately 3 Å in the TAPBPR-bound complex relative to the fully loaded pMHC-I structure (measured at the Cα atom of A150, **Figure 4B**), in line with the observations made on the basis of empty MHC-I/TAPBPR complexes^28,29^. Notably, in contrast to the previous empty TAPBPR/MHC-I crystal structures as well as the fully loaded pMHC-I structure, our cryoEM structure reveals a significant alteration of the α_1_ helix and adjacent loop (residues S71-G91) (**Figure 4B, Movie S2**), where a lack of clear electron density indicates the presence of conformational heterogeneity impacting the F-pocket of the peptide-binding groove (**Figure S8**). This finding reflects the dynamic transitions of the F-pocket between peptide-deficient and peptide-bound states in a solution environment (**movie S2**), in agreement with our previous solution-based NMR studies of MHC-I/TAPBPR complexes^32^.

**Figure 4.**
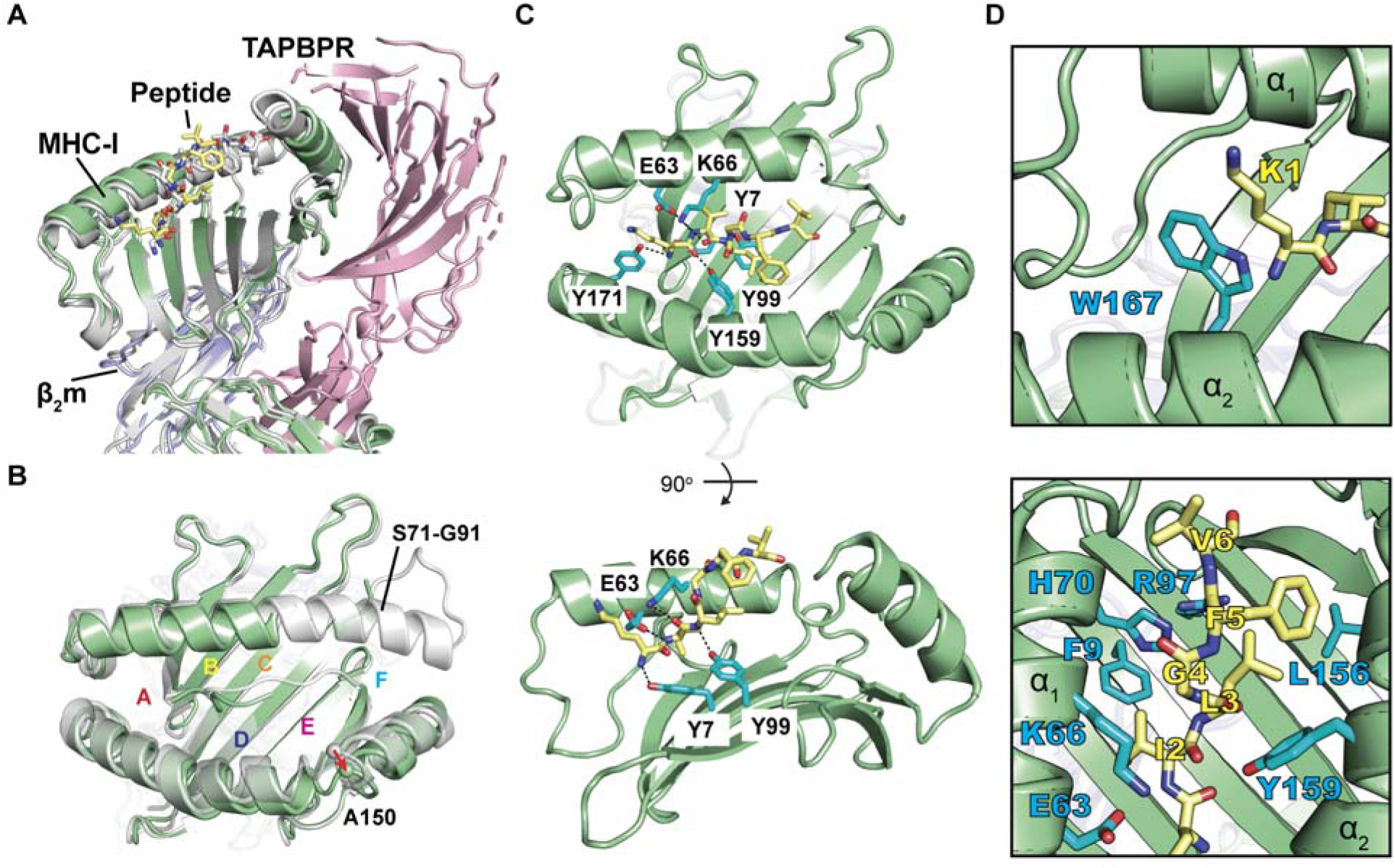
Structural changes between free and chaperoned peptide-bound HLA-A*02:01. (**A**) Overview of the molecular complex between TAPBPR^HiFi^/HLA-A*02:01/β_2_m with a peptide decoy (PDB ID 9C96, EMDB ID EMD-45360), shown as light pink, pale green, light blue cartoons, and yellow sticks, respectively, overlaying with peptide-loaded (GILGFVFTL) HLA-A*02:01 (PDB ID 2VLL) shown in gray. The MHC-I α_2_helix has been partially removed for peptide visualization. (**B**) Structural superposition of chaperone-free peptide-loaded HLA-A*02:01 (PDB ID 2VLL) onto the cryoEM structure of chaperoned KILGFVF/HLA-A*02:01 (PDB ID 9C96, EMDB ID EMD-45360) showing well-resolved density for peptide residues 1-6, and a lack of cryoEM density for residues S71-G91 of HLA-A*02:01. MHC-I A-, B-, C-, D-, E-, and F-pockets are noted and the spheres are showing A150 Cα. (**C**) Detailed interactions between MHC-I groove residues with the backbone of peptide residues P1-P3, as indicated (top and side views). Hydrogen bonds are indicated by black dashed lines. (**D**) Close-up view of the molecular interactions between the N-terminal peptide hydrophobic sidechain with MHC-I peptide binding groove residues. See also **Figure S9**.

Compared to the original influenza epitope GILGFVFTL, the heptamer peptide decoy (KILGFVF) within the TAPBPR editing complex exhibits a native-like backbone conformation and sidechain rotamer placement (**Figure 4A and C, Movie S1**). Despite the highly polymorphic character of the MHC-I peptide binding groove, a relatively conserved cluster of residues (Y7, E63, K66, Y99, Y159, and Y171) located within the A-, B-, and D-pockets mediate stable docking interactions with the peptide backbone (**Figure 4C and S9, Movie S1**)^47^. Additional groove residues, including W167, F9, H70, L156, and Y159, with hydrophobic sidechain anchors the N-terminus of the peptide, particularly at the P2 and P3 positions (**Figure 4D, Movie S1**). While some amino acid polymorphisms across common HLA allotypes can be observed at positions 63 and 66 (E63 and K66 for HLA-A*02:01), with the exception of I66, all other amino acid substitutions have the potential to serve as hydrogen bond donors to the carbonyl oxygen of the peptide at P3 (**Figure 4D**). However, the dynamic character of the S71-G91 region in the editing complex leads to missing interactions between the C-terminus of the peptide with residues D77, Y84, T143, and W147 of the E- and F-pockets (**Figure S9, Movie S1**). The flexibility of the F pocket serves as an additional filter to screen for high-affinity peptides with the desired C terminal amino acids.

Placing in the context of previous NMR and X-ray studies^21,28,29,32,41^, our cryo-EM structure demonstrates that the initial capture of peptides for proofreading by TAPBPR occurs through native-like interactions with conserved MHC-I residues within the A-, B-, and D-pockets that are properly conformed to receive peptide. The F-pocket is partially formed in this intermediate state of the complex and folds upon annealing of peptide C-terminus for high-affinity ligands with robust P9 anchors, allosterically shifting the groove inward via α_2-1_ helix and promoting the dissociation of TAPBPR from the pMHC-I complex^32^. These functional and structural results altogether suggest that peptide-loaded MHC-I can form high-affinity, stable proofreading complexes with TAPBPR^HiFi^ for antigen selection.

### TAPBPR^HiFi^ enhances loading of exogenous antigens on the cell surface

We hypothesized that TAPBPR^HiFi^ could enhance the exchange of peptides on different HLA-A allotypes expressed on a cellular surface. Using a flow cytometry-based method, we assessed the loading of fluorophore-labeled peptides upon incubation with soluble TAPBPR variants, TAPBPR^HiFi^ TAPBPR^WT^, and TAPBPR^TN6^, at different concentrations for monoallelic 722.211 cell lines^48^ expressing either HLA-A*02:01, 24:02, 23:01, or 03:01 (**Figure 5A and S10**). Consistent with our bead-based binding data, we found that a nanomolar-range concentration of TAPBPR^HiFi^ can readily exchange peptides on HLA-A*02:01, relative to the negative control TAPBPR^TN6^, whereas TAPBPR^WT^ shows no significant enhancement in exchanging for targeted fluorophore-labeled peptides (**Figure 5B and S11A**). Similarly, using monoallelic HLA-A*24:02 and HLA-A*23:01 cell lines, we observed that TAPBPR^HiFi^ can promote robust peptide exchange at a concentration that is two orders of magnitude lower than TAPBPR^WT^ (**Figure 5B and S11B-C**). While showing lower levels of binding to TAPBPR PE-tetramers in our SAB assay, nanomolar soluble TAPBPR^HiFi^ can promote loading of exogenous antigens for HLA-A*03:01, albeit the use of a higher peptide concentration relative to the other HLA allotypes (**Figure 5B and S11D**). We also evaluated whether TAPBPR^HiFi^-TM^18^ expressed on the cell surface of TAP transporter deficient T2 cells that lack the endogenous antigen processing machinery can directly enhance the loading of exogenous, fluorescently labeled peptides (**Figure 5C and S12A**). Our results showed that TAPBPR^HiFi^ can significantly enhance the level of peptide loading at low peptide concentrations relative to T2 cells expressing TAPBPR^TN6^ and TAPBPR^WT^-TM (**Figure 5D and S12C-D**). We further demonstrate that surface-expressed TAPBPR^HiFi^-TM increases the expression level of HLA-A*02:01 on the T2 cell surface, as indicated by staining with the allele-specific BB7.2 anti-HLA A2 antibody PE (**Figure S12D**). HLA-A*02:01 molecules on T2 cells expressing TAPBPR^HiFi^-TM are receptive to peptide binding at a picomolar peptide concentration (**Figure 5D and S12D**). Taken together, our results demonstrate that our engineered high-fidelity TAPBPR variant TAPBPR^HiFi^ can have a wide range of applications in promoting peptide exchange and participating in antigen selection across different HLA-A allotypes on the cell surface.

**Figure 5.**
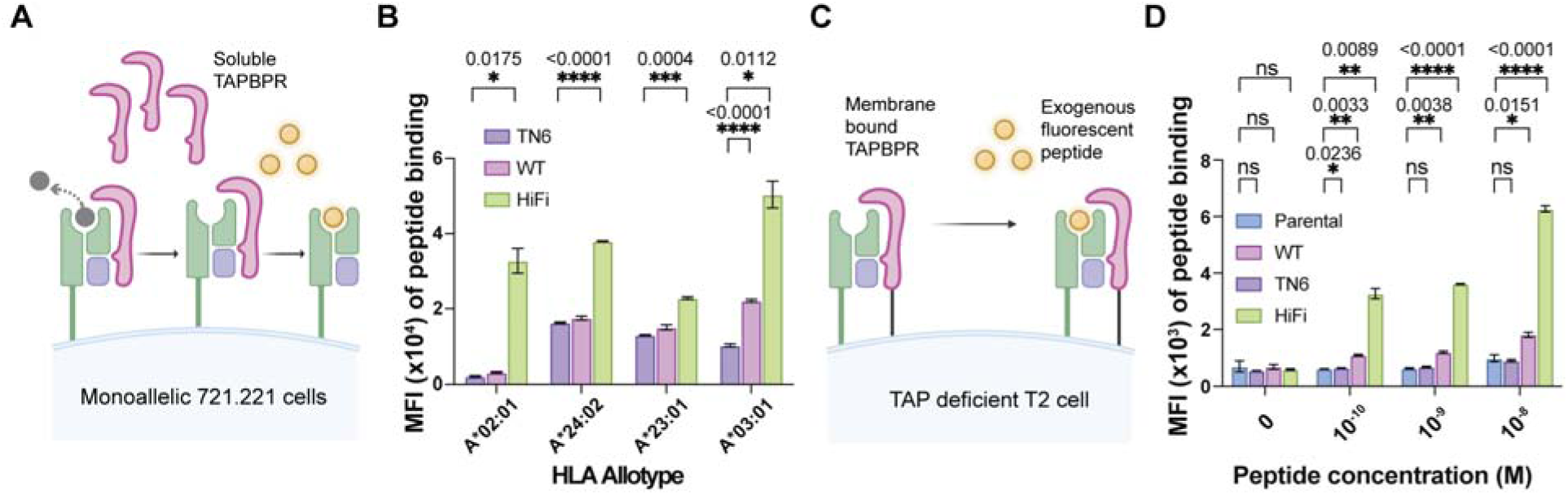
TAPBPR^HiFi^ enhances peptide loading across common HLA-A* allotypes expressed on a cellular membrane. (**A**) Schematic of peptide exchange for MHC-I expressed on the cell surface by soluble TAPBPR. (**B**) Bar graph summarizing the median fluorescence intensity (MFI) of fluorescent peptide binding for monoallelic HLA-A*02:01, A*24:02, and A23:01 cell lines with 10 nM fluorescent peptide and 1 μM TAPBPR^HiFi^, TAPBPR^WT^, or TAPBPR^TN6^, and HLA-A*03:01 cell line in the presence of 10 μM fluorescent peptide and 10 μM TAPBPR^HiFi^, TAPBPR^WT^, or TAPBPR^TN6^, as indicated, from three independent experiments. Two-way ANOVA was performed relative to TAPBPR^TN6^, P > 0.0.1234 (ns), P < 0.0332(*), P < 0.0021(**), P < 0.0002(***), and P < 0.0001(****). (**C**) Schematic of peptide exchange for MHC-I expressed on the cell surface by membrane-bound TAPBPR. (**D**) Bar graph summarizing the MFI of fluorescent peptide binding to HLA-A*02:01 expressed T2 cell line with no transduction (parental) or TAPBPR^HiFi^, TAPBPR^WT^, or TAPBPR^TN6^-TM transduction, in the presence of different concentrations of fluorophore-labeled peptide as indicated, from three independent experiments. Two-way ANOVA was performed relative to the parental (no transduction), P > 0.0.1234 (ns), P < 0.0332(*), P < 0.0021(**), P < 0.0002(***), and P < 0.0001(****). See also **Figure S10-S12**.

**Figure 6.**
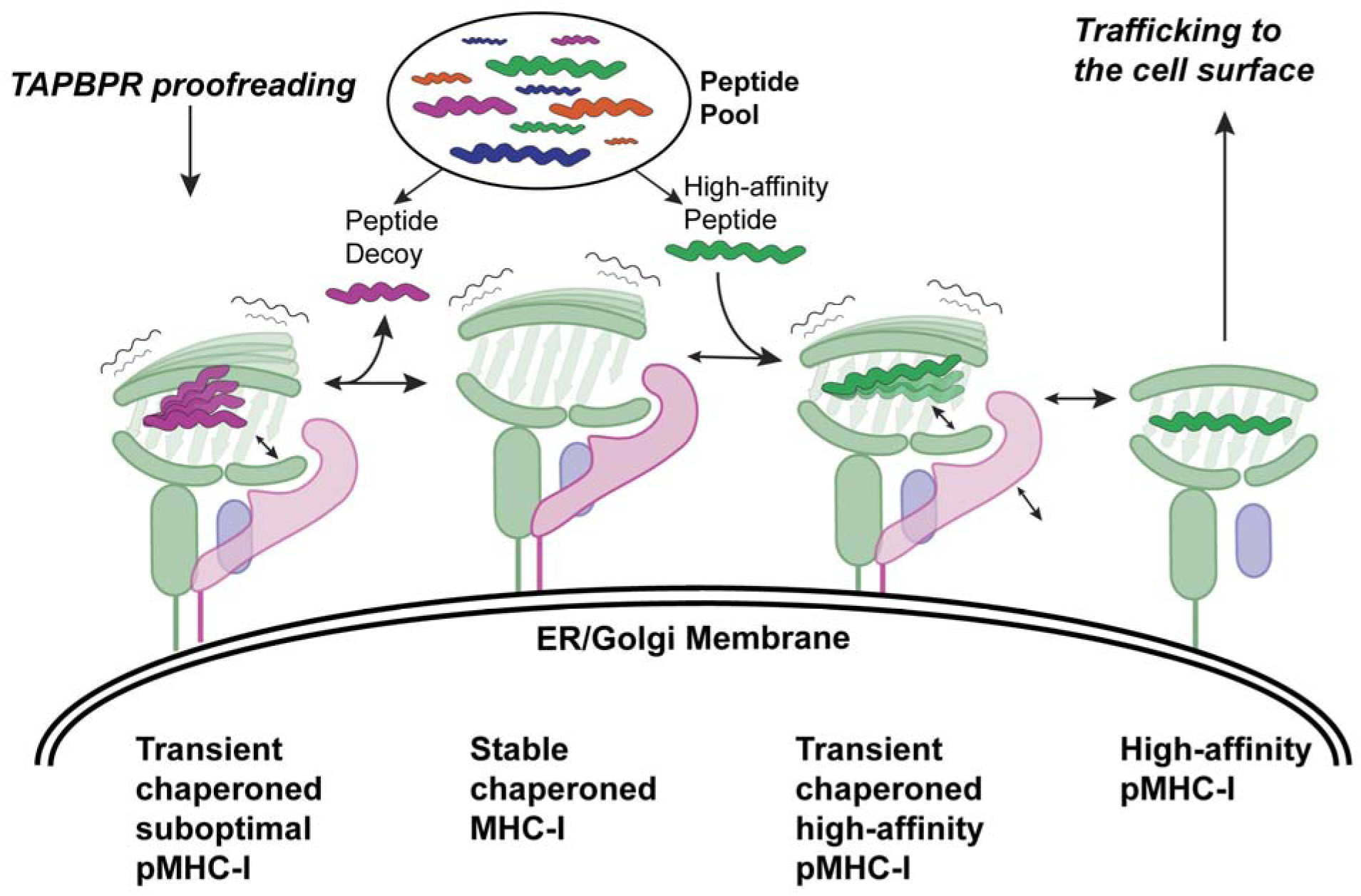
Structural mechanism of peptide antigen proofreading by TAPBPR. Empty MHC-I is preferentially recognized by TAPBPR in a peptide-receptive, open conformation. Chaperoned MHC-I screens a large peptide pool in the endoplasmic reticulum through transient complexes. TABBPR widens the α_2-1_ helix of the peptide binding groove, enhances the dynamics of the α_1_ helix, and induces an unstructured F-pocket. Thus, chaperone MHC-I exhibits a lower affinity towards incoming peptides and promotes the dissociation of suboptimal peptide decoys. When chaperoned MHC-I proofreads a high-affinity peptide, the N terminal peptide forms native-like contacts with its A- and B-pockets. The peptide C-terminus interacts with peptide-coordinating MHC-I residues in the F-pocket to stabilize the peptide binding groove in a closed conformation, which allosterically promotes the release of TAPBPR.

## Discussion

The molecular chaperones tapasin and TAPBPR play important roles in stabilizing nascent MHC-I molecules, optimizing the repertoire of bound peptide cargo, and mediating quality control of peptide-loaded molecules, in tandem with other components of the MHC-I antigen processing pathway^1,2,49–51^. Advances in sample preparation and the use of complementary structural techniques have provided a range of resting-state (empty) MHC-I/chaperone complex structures, which have established the basis of tapasin and TAPBPR recognition of peptide-deficient MHC-I molecules^52,53^. Notwithstanding, the molecular mechanism of peptide editing is enigmatic and highly controversial in the literature^36^. In previous work, we used solution NMR methyl probes to study a peptide-bound MHC-I/TAPBPR intermediate and showed that it involves a transient conformational state (approximately 200 millisecond lifetime)^32^. A large body of work has established that dynamics at residues distributed throughout the MHC-I peptide binding groove are dampened upon binding of high-affinity peptides, and allosterically coupled to the TAPBPR binding surfaces on the underside and α_2-1_helix of the groove to promote chaperone release from the pMHC-I ^21,32,41^. Recent X-ray structures of HLA-B*08:01 bound to 20mer peptides with protruding N-terminus have provided clues into the peptide-bound intermediate of the loading process^54^. Despite these insights, a high-resolution structure of the transient peptide-bound chaperoned MHC-I has been missing due to its transient nature and dynamic complexity, hindering structure determination by X-ray crystallography or cryoEM.

We here leverage recent advances in protein engineering of the individual components, a high-fidelity TAPBPR^HiFi^ variant^18^, and an ultra-stable, open HLA-A*02:01 molecule^38^ prepared using a photocleavable conditional peptide ligand^55^, to isolate a TAPBPR/MHC-I intermediate bound to a heptamer peptide decoy. The insights gained from our peptide-loaded MHC-I/TAPBPR cryoEM structure allow us to propose the following mechanism for peptide proofreading on MHC-I (Fig. 5). Empty MHC-I is preferentially recognized and stabilized by TAPBPR in a peptide-receptive, “open” state. Chaperoned MHC-I molecules screen the large peptide pool within the endoplasmic reticulum compartment, transiently forming the peptide/MHC-I/TAPBPR complex. Meanwhile, TABBPR widens the peptide binding groove via the α_2-1_ helix and enhances the dynamic movements alongside the α_1_helix and F pocket. As a result, the key peptide-coordinating residues within the E- and F-pockets lose their interactions with the peptide C-terminus. Therefore, TAPBPR-bound MHC-I exhibits an overall lower affinity towards incoming peptides and promotes the dissociation of transiently bound peptide decoys. When chaperoned MHC-I proofreads a high-affinity peptide, it interacts with the N terminal peptide through native-like contacts formed within the A-, B-, and D-pockets. The C-terminus of the high-affinity peptide interacts with peptide-coordinating residues in the E- and F-pockets to stabilize and close the peptide binding groove, which allosterically triggers the release of TAPBPR^32^. The resulting high-affinity peptide-loaded MHC-I that has been proofread by TAPBPR can traffic to the cell surface for antigen presentation.

The TAPBPR G24-R36 loop has been previously suggested to directly mediate peptide editing by competing with the peptide C-terminus for binding to the MHC-I F-pocket^29,39,40^. More recent studies using solution NMR have challenged this “scooping/levering” mechanism by explicitly showing that the TAPBPR loop adopts a disordered conformation, which forms hydrophobic contacts with the rim of the α_2-1_ helix, instead of entering the empty MHC-I groove in a solution environment^39–41^. Our engineered high-affinity TAPBPR^HiFi^ corroborates this model, by showing that the S104F mutation on the edge of the TAPBPR loop contributes to enhanced binding on peptide-loaded MHC-I molecules. In further agreement with our previous NMR studies^41^ and the lack of clear electron density in the structure of both MHC-I/TAPBPR complexes^28,29^, we find that, even though the F-pocket of the MHC-I groove in complex with the heptamer peptide decoy is completely empty, electron density for the loop residues G24-R36 is completely lacking in our data, supporting a more disordered ensemble of conformations. Taken together, our results support that TAPBPR promotes peptide editing by distorting the regular structure of the MHC-I F-pocket, including a widened α_2-1_ and a partially unfolded α_1_ helix, rather than by directly competing with the peptide terminus. These insights provide a complete view of how TAPBPR proofreads the intracellular peptide pool to select an optimized repertoire of epitopes for MHC-I antigen presentation on the cell surface.

## Acknowledgments

The authors acknowledge Dr. Hoang-Anh Phan (CHOP) for assistance with mass spectrometry and Sagar Gupta for assistance with preparing movies and peptide sequence logos. We are grateful to Dr. Derin Keskin (Harvard University) for sharing the HLA monoallelic cell lines, and Dr. Frank DiMaio (University of Washington) for assistance with structure modeling using Rosetta. We also acknowledge the use of instruments at the Electron Microscopy Resource Lab and at the Beckman Center for Cryo-Electron Microscopy at the Institute of Structural Biology at the University of Pennsylvania Perelman School of Medicine. We thank Prerana Gogoi and Sudheer Molugu for assistance with Krios microscope operation at the University of Pennsylvania.

## Funding

NIH grants R01AI143997 (N.G.S.)

R35GM125034 (N.G.S.)

R35GM144120 (V.M.B)

The Children’s Hospital of Philadelphia Cell and Gene Therapy Collaborative

## Contributions

Conceptualization: NGS.

Methodology: YS, RAP, LM, CW, DH, AC, JND, KD, VM-B

Investigation: YS, RAP, LM, CW, DH, AC, JND, KD

Visualization: YS, RAP

Funding acquisition: NGS, VM-B

Supervision: NGS

Writing – original draft: NGS, YS, RAP,

Writing – review & editing: NGS, VM-B, YS, LM, CW, DH, AC, JND, KDG

## Data and materials availability

All structures and cryoEM density data associated with this study have been deposited in the PDB and EMDB, under PDB ID 9C96 and EMDB entry ID EMD-45360. NMR assignments have been deposited in the BMRB, under code 52520. All other data relating to this paper are deposited in Dryad via doi:10.5061/dryad.prr4xgxvv. (https://datadryad.org/stash/share/U_rx4_y77uxIogWeqEldx4uY_etUhheCgH2FTWN7WIw can be accessed by the reviewers)

