## Supplementary Information for "CryoEM structure of an MHC-I/TAPBPR peptide bound intermediate reveals the mechanism of antigen proofreading"

**The PDF file includes:**

Materials and Methods

References

Figures S1 to S12

**Other Supplementary Materials for this manuscript include the following:**

Tables S1 to S2

Movies S1 to S2

Data S1

Materials and Methods

Peptides

All peptide sequences are given as standard single-letter codes. High-affinity peptide TAX9 (LLFGYPVYV) was purchased from Genscript, USA, at >90% purity. The fluorophore-labeled and photolabile peptides, using TAMRA as 5-Carboxytetramethylrhodamine and J as Fmoc-3-amino-3-(2-nitrophenyl)-propionic acid, were purchased from Biopeptek Inc, Malvern, USA. Peptides were solubilized in distilled water and centrifuged at 14000 rpm for 15 minutes. The concentration of each peptide solution was measured and calculated using the absorbance and extinction coefficient at 205 nm wavelength.

Recombinant protein expression, refolding, and purification

DNA Plasmids encoding the luminal domain of MHC-I heavy chain HLA-A*02:01 and light chain human β_2_m were provided by the NIH tetramer facility (Emory University). Plasmids encoding open HLA-A*02:01 (G120C) and open β_2_m (G120C) were site mutated in-house. Paired plasmid were individually transformed into Escherichia coli BL21 (DE3) cells (New England Biolabs). Proteins were expressed in the Luria-Broth medium, and inclusion bodies were collected and purified as previously described^1^. For the generation of peptide-loaded HLA-A*02:01 (pHLA-A*02:01) molecules, in vitro refolding was performed by slowly diluting a 200 mg mixture of MHC-I heavy chain and β_2_m at a 1:3 molar ratio over 24 hours in refolding buffer (0.4 M L-Arginine HCl, 100 mM Tris pH 8, 2 mM EDTA, 4.9 mM reduced L-glutathione, 0.57 mM oxidized L-glutathione) containing 10 mg of the desired peptide. The mixture was protected from light when refolded with photolabile peptides. Refolding proceeded for four days, and proteins were purified by size exclusion chromatography (SEC) using a HiLoad 16/600 Superdex 75 pg column at 1 mL/min with 150 mM NaCl, 20 mM Tris buffer, pH 8.0.

The luminal domain of TAPBPR and its variant proteins containing the non-disulfide bonded free cysteine mutation (C94S) tagged with BSP and 6-His was stably expressed in the Drosophila melanogaster S2 cell line^2^. The cultures were induced with 1 mM CuSO4, and the supernatant was collected after four days. The secreted TAPBPR molecules were purified using a high-density metal affinity agarose resin (ABT, Madrid). The eluted proteins were further purified by SEC using a HiLoad 16/600 Superdex 200 pg column at a 1 mL/min flow rate in 150 mM NaCl, 20 mM sodium phosphate buffer, pH 7.4.

Differential scanning fluorimetry

DSF was used to measure the thermal stability of the TAPBPR proteins. TAPBPR proteins (7 μM) were mixed with 10X SYPRO Orange dye in a buffer of 150 mM NaCl and 20 mM sodium phosphate (pH 7.2) to a final volume of 20 μl. Samples were loaded into a MicroAmp Optical 384-well plate and ran in triplicates. The experiment was performed on a QuantStudio 5 real-time polymerase chain reaction (PCR) machine with excitation and emission wavelengths set to 470 and 569nm. The thermal stabilities of TAPBPR were generated by plotting the first derivative of each melting curve and extracting the peak as the melting temperature (Tm). The thermal stability was measured by gradually increasing temperature at a rate of 1°C/min between 25° and 95°C. Data analysis and fitting were performed in GraphPad Prism v10.

Surface plasmon resonance

SPR experiments were conducted in triplicate using a Biacore X100 instrument (Cytiva) in SPR buffer (150 mM NaCl, 20 mM sodium phosphate pH 7.4, 0.1% Tween-20). Approximately 1000 resonance units (RU) of biotinylated TAX9/HLA-A*02:01 were immobilized at 10 μL/min on a streptavidin-coated chip (Cytiva). The chip was primed by flowing over 1 μM TAX9 peptide solution before each binding assay to prevent peptide release from the pMHC-I molecules. Various concentrations from 0.1 up to 300 μM of TAPBPR and its variants were injected onto the chip at 25˚C at a flow rate of 30 μL/min for 60 seconds, followed by 180s dissociation time. The SPR sensorgrams and equilibrium dissociation constants (K_D_) were analyzed using the surface-bound analysis settings in Biacore X100 evaluation software (Cytiva). The representative SPR sensorgrams and fitted saturation curves were prepared in GraphPad Prism v9. The K_D_s of TAPBPR^WT^, TAPBPR^∆ALAS^, and TAPBPR^∆G24-R36^ are estimated values since saturation of binding was not reached.

Fluorescence polarization

The fluorescent peptide dissociation of the fluorophore-labeled peptide-loaded MHC-I was monitored by fluorescence polarization (FP). TAMRA-TAX9/HLA-A*02:01 protein at a concentration of 40 nM in the presence of 1 μM non-labeled TAX9 peptide was incubated without chaperone or with 1 μM TAPBPR^TN6^, TAPBPR^WT^, TAPBPR^∆G24-R36^, TAPBPR^HiFi^, and TAPBPR ^HiFi∆G24-R36^. The kinetic dissociations were monitored for 10 hours, and fluorescence polarization measurements were recorded approximately every 90 seconds. Excitation and emission values used to monitor the fluorescence of TAMRA-labeled peptides were 531 and 595 nm. All experiments were performed at RT in triplicates. Raw parallel (I_II_) and perpendicular emission intensities (I⊥) were collected and converted to polarization (mP) values using the equation 1000*[(I_II_-(G*I_⊥_))/(I_II_+(G*I_⊥_))]. An optimized G-factor was determined to be 0.33 for TAMRA-labeled peptides in calculating baseline fluorescence and overall fluorescence polarization. The plateau for each dissociation was extracted by fitting one phase decay in GraphPad Prism 10. The relative dissociation was calculated using the equation $\% relative dissociation= \frac{\mathrm{Plateau}^{buffer only}-\mathrm{Plateau}^{\mathrm{chaperone}}}{\mathrm{Plateau}^{buffer only}}$. Error bar (SD) was propagated from three independent experiments.

Single antigen bead (SAB) screen

TAPBPR^HiFi^, TAPBPR^WT^, and TAPBPR^TN6^ (7 μM) phycoerythrin (PE)-tetramers were mixed with 4 μl of LABScreen SAB suspension (OneLambda Inc., CA, USA) in a 96-well plate. The samples were incubated for 1 hour and 550 rpm at RT, washed four times in wash buffer (OneLambda Inc., CA, USA) to remove excess tetramers, and resuspended in phosphate-buffered saline (PBS; pH 7.2). For the negative controls, beads were incubated with the anti–HLA class I antibody W6/32 (Abcam, ab22432) at 550 rpm for 30 minutes at room temperature and washed three times, followed by TAPBPR PE-tetramer addition. We measured the mean fluorescent intensity (MFI) of SABs upon incubation with TAPBPR PE-tetramers. We then used the MFI of corresponding TAPBPR PE-tetramer staining of SABs that are pre-incubated with W6/32 as the negative control. The MFI ratio was then calculated using the MFI of TAPBPR staining divided by the MFI of the corresponding negative control staining^3^. To test the levels of peptide-loaded MHC-I molecules on the beads, we used the same W6/32 antibody and the secondary anti-mouse PE-conjugated antibody (Abcam, ab97024) for detection. The levels of TAPBPR bound to the beads were measured using the Luminex 100 Liquid Array Analyzer System, and the results were analyzed in GraphPad Prism v10.

Fluorescent peptide exchange on a cellular membrane

721.221 Human HLA negative B-lymphoblastoid cells and monoallelic HLA-A*02:01, 24:02, 23:01, and 03:01 722.211 cell lines^4^ were thawed and expanded in RPMI complete medium (RPMI-1640 with 25 mM HEPES & L-Glutamine, 10% heat-inactivated FBS, 1 mM sodium pyruvate, and 1% pen-strep). These cell lines were maintained at a density of 200,000 – 250,000 cells/mL. Cells were resuspended to a concentration of 2X10^6^ cells/mL and plate 200,000 per well and then stained with LIVE/DEAD™ Fixable Violet for 10 minutes at RT and for 30 minutes at 4 °C in 2% BSA containing one of the following fluorophore-labeled antibodies: anti-HLA A2 antibody PE, anti-HLA A24 antibody FITC, anti-HLA A3 antibody APC. Cells were washed three rounds to remove excess unbound fluorophore-labeled antibodies The acquisition was then performed on CytoFLEX LX (Beckman Coulter), and the data were analyzed by FlowJo v10.10.0.

Upon validating the surface expression of HLA-A*02:01, 24:02, 23:01, and 03:01, target cell lines were seeded at 200,000 cells per well for peptide exchange. These cells were then treated with recombinant TAPBPR^HiFi^, TAPBPR^WT^, or TAPBPR^TN6^ at different concentrations, as indicated, for 15 minutes at 37 °C. After 15 minutes, the corresponding fluorophore-labeled peptide, 10nM TAMRA-TAX9 (KLFGYPVYV) for HLA-A*02:01, 10nM TAMRA-PHOX2B (KYNPIRTTF) for HLA-24:02 and 23:01, or 10 μM FITC-PLP (KLIETYFSK, proteolipid protein epitope) for HLA-A*03:01, was added to the cells and incubated at 37 for 60 minutes. Following the peptide treatment, the cells were washed three times in 1xPBS and harvested. The acquisition was performed on CytoFLEX LX (Beckman Coulter) using the PE or FITC channel, and the data were analyzed by FlowJo v10.10.0.

Lentiviral production, transduction, and peptide exchange on TAPBPR-TM transduced cells

Three plasmids were used for lentivirus production: pMD2.G (envelope plasmid encoding VSV-G), psPAX2 (packaging plasmid), and pSFFV (transfer plasmid containing FLAG-tagged TAPBPR variants). Lenti-X 293 cells were cultured in Dulbecco’s Modified Eagle’s Medium (DMEM) supplemented with 10% fetal bovine serum (FBS). For lentivirus production, Lenti-X 293 cells were grown to 70-80% confluence in a T75 flask and were co-transfected with psPAX2, pSFFV, and pMD2.G using Lipofectamine 3000. Lentivirus containing supernatant was collected every 24 hours for 3 total harvests and concentrated using Lenti-X concentrator as suggested by the manufacturer (Takara). The viral pellet was resuspended in phosphate-buffered saline (PBS), flash-frozen in liquid nitrogen, and stored at -80°C for further use. Transductions were performed using retronectin-coated plates (Takara) and different titers of lentivirus along with 1x10^5^ T2 cells in a 96-well plate. Cells were split every 1-2 days for up to 4 days before analysis by flow cytometry for TAPBPR expression for TAPBPR^HiFi^, TAPBPR^WT^, and TAPBPR^TN6^-TM transduced cell lines. Cells with similar levels of transduction efficiency and TAPBPR expression were selected for downstream assays, as measured by staining for FLAG (Clone: L5 in APC). Expression levels of cell surface HLA-A*02:01 were quantified by staining with an anti-HLA A2 antibody (Clone: BB7.2 in Brilliant Violet 785). T2 cells transduced with TAPBPR variants were incubated with indicated fluorescent TAX9-TAMRA peptide conjugate concentrations for 60 minutes in serum-free DMEM. Cells were then washed with FACS buffer, stained with LIVE/DEAD Fixable Near IR 876 (Invitrogen) for viability, and fixed with 4% PFA in 1x PBS. Cells were subsequently analyzed by flow cytometry using a CytoFlex LX cytometer.

Peptide-loaded open HLA-A*02:01/TAPBPR^HiFi^ complex purification

Open KILGFVFJV/HLA-A*02:01/β_2_m developed with enhanced stability in our previous work^5^ was mixed with TAPBPR^HiFi^ at a 1:1.3 molar ratio. The mixture was UV-irradiated at a high intensity in 150 mM NaCl, 20 mM sodium phosphate, pH 7.4 at 4˚C for 40 minutes with a wavelength of 365 nm. The complex was then purified by SEC using a Superdex 200 pg Increase 10/300 GL, and the eluted peaks were further analyzed by Sodium Dodecyl Sulphate/ Polyacrylamide Gel Electrophoresis (SDS/PAGE) to identify all components and confirm the complex. The open HLA-A*02:01/TAPBPR^HiFi^ complex was then prepared at a 0.2 mg/mL concentration with 1XPBS, pH 7.4 for cryoEM screen.

NMR Titrations and Chemical Shift Mapping.

NMR chemical shift mapping for the TAX9/HLA-A*02:01 molecule upon binding to TAPBPR^HiFi^ was performed in an analogous fashion to TAPBPR^WT^, described in our previous work^6^. ^13^CH_3_ AILV-labeled HLA-A*02:01 at 103 μM was titrated with increasing concentrations of unlabeled TAPBPR^HiFi^ in matched NMR buffer (20 mM sodium phosphate, 100 mM NaCl, 10% D2O (v/v) at pH 7.2). The HLA-A*02:01: TAPBPR^HiFi^ ratios were: 1: 0, 1: 0.55, 1: 1.10, 1: 1.65, 1: 2.75, and 1: 3.31. Two-dimensional methyl ^1^H-^13^C methyl SOFAST HMQC experiments were recorded at 298 K at a field strength of 600 MHz with a total number of 136 scans, a recycle delay (d1) of 0.2 s and an acquisition time of 30 milliseconds in the indirect dimension. Chemical shift deviations (CSD) were computed using the equation Δδ_CH3_ = [ ½ (Δδ_1H_^2^ + Δδ_13C_^2^/4)] ^½^ for the AILV methyl groups, where Δδ_1H_ and Δδ_13C_ are the changes in proton and carbon chemical shifts, respectively, upon saturation with TAPBPR^HiFi^. Using the TITAN software, the dissociation constant was obtained by fitting the slow-exchanging NMR resonances^7^. For this analysis, the titration spectra were processed using an exponential window function with 4 Hz and 10 Hz line broadening in the direct and indirect dimensions, respectively, and fit using a two-state binding model in TITAN with bootstrap error analysis of 100 replicas.

Liquid chromatography-mass spectroscopy (LC-MS)

LC-MS was carried out with the passage of protein complex or peptide solution at a concentration range of 5-10 μM a Waters Acquity C8 column followed by electron ion spray–MS performed on a Waters Acquity UPLC SQD instrument with a mass detection range of 0-2000 m/z. Analysis and deconvolution of LC-MS data were performed with Xcalibur and UniDec.

CryoEM sample preparation and data collection

The CryoEM sample was prepared as previously described. Briefly, the open peptide-loaded HLA-A*02:01/TAPBPR^HiFi^ complex was obtained by mixing open HLA-A*02:02/TAX9 and TAPBPR^HiFi^ at a 1:1.3 molar ratio. The mixture was incubated at 4 ˚C for 1 hour and followed by 40-minute UV irradiation, and the complex was purified by size-exclusion chromatography. The peak fraction at 0.2 mg/mL concentration was used for grid preparation. The sample was applied to freshly plasma cleaned Quantifoil Cu 300 2/2 (Quantifoil) grids and was plunge frozen in liquid ethane using the Vitroblot Mark IV (Thermo Fisher) operated at 4°C and 100% humidity.

The dataset was collected on a Titan Krios G3i 300 kV electron microscope (Thermo Fisher Scientific) with a 20 eV energy filter and equipped with a K3 Summit camera (Gatan). Super-resolution images were collected over thirty-five frames with a dose of 40.5 e-/A^2^ at a nominal magnification of x105,000, resulting in a pixel size of 0.418 Å/pixel. The defocus range was set from -0.8 to -3.0 μm.

CryoEM Data Processing

CryoEM data processing was performed using Relion 5.0^8^ and cryoSPARC 4.5^9^. 5,381 collected movies were motion corrected using the Relion implementation of MotionCor2^10^, binning to a pixel size of 0.836 Å/pixel. The defocus values were determined using CTFFIND-4^11^ and then suboptimal micrographs were removed, resulting in a set of 4,791 micrographs. From this point, processing proceeded in cryoSPARC. Templates for particle picking were generated using blob picker and 2D classification on a subset of the dataset, which was also used to generate a good ab initio model. Template picking resulted in 4,707,523 particles, which were extracted and binned to a pixel size of 3.344 Å/pix. These particles were split evenly into five subsets and each subset was subjected to hetero refinement, using a single copy of the good ab initio model and five copies of a noise volume. After this first round of hetero refinement, a collective 1,119,002 particles were sorted into the good class. The 1,119,002 particles were combined and again split into two even subsets and subjected to the same hetero refinement protocol again, resulting in 628,062 particles sorted into the good class. The 628,062 particles were then combined and re-extracted to a pixel size of 1.672 Å/pixels, and any overlapping particles were removed. These particles were then subjected to eight additional rounds of the same hetero refinement protocol, until particle loss to the bad classes became negligible, resulting in a set of 340,226 particles. Throughout this process, particles from both the good class and the bad classes were subjected to 2D classification as a sanity check to ensure good particles were not discarded. These 340,226 particles were re-extracted to their final box size of 0.836 A/pixel and subjected to non-uniform refinement^12^, resulting in a 3.3 Å structure. These particles were then transferred to Relion and reconnected to the original motion corrected micrographs. The 340,226 particles were refined in Relion using Blush regularization^13^, resulting in a 3.3 Å structure. The use of Blush regularization significantly improved the map quality compared to cryoSPARC refinement or Relion refinement without Blush regularization. The particles were then subjected to two rounds of CTF refinement^14^ and Bayesian polishing, resulting in a 3.0 Å structure. As significant heterogeneity for the peptide was still present, the 340,226 particles were then locally refined centered on the MHC peptide groove, followed by classification without angular assignment of the same region. This resulted in a set of 88,714 particles with the best density for the peptide in the MHC groove. These particles were then subjected to an overall refinement with Blush regularization, resulting in a 3.0 Å structure after postprocessing in Relion.

Model building and structure refinement

The model for the MHC-I/TAPBPR complex was built iteratively using ISOLDE^15^, Coot^16^, and the PHENIX^17^ software package, using PDB 2VLL as the starting model. Images of the models and maps for figures were generated using ChimeraX^18^ and Pymol^19^.

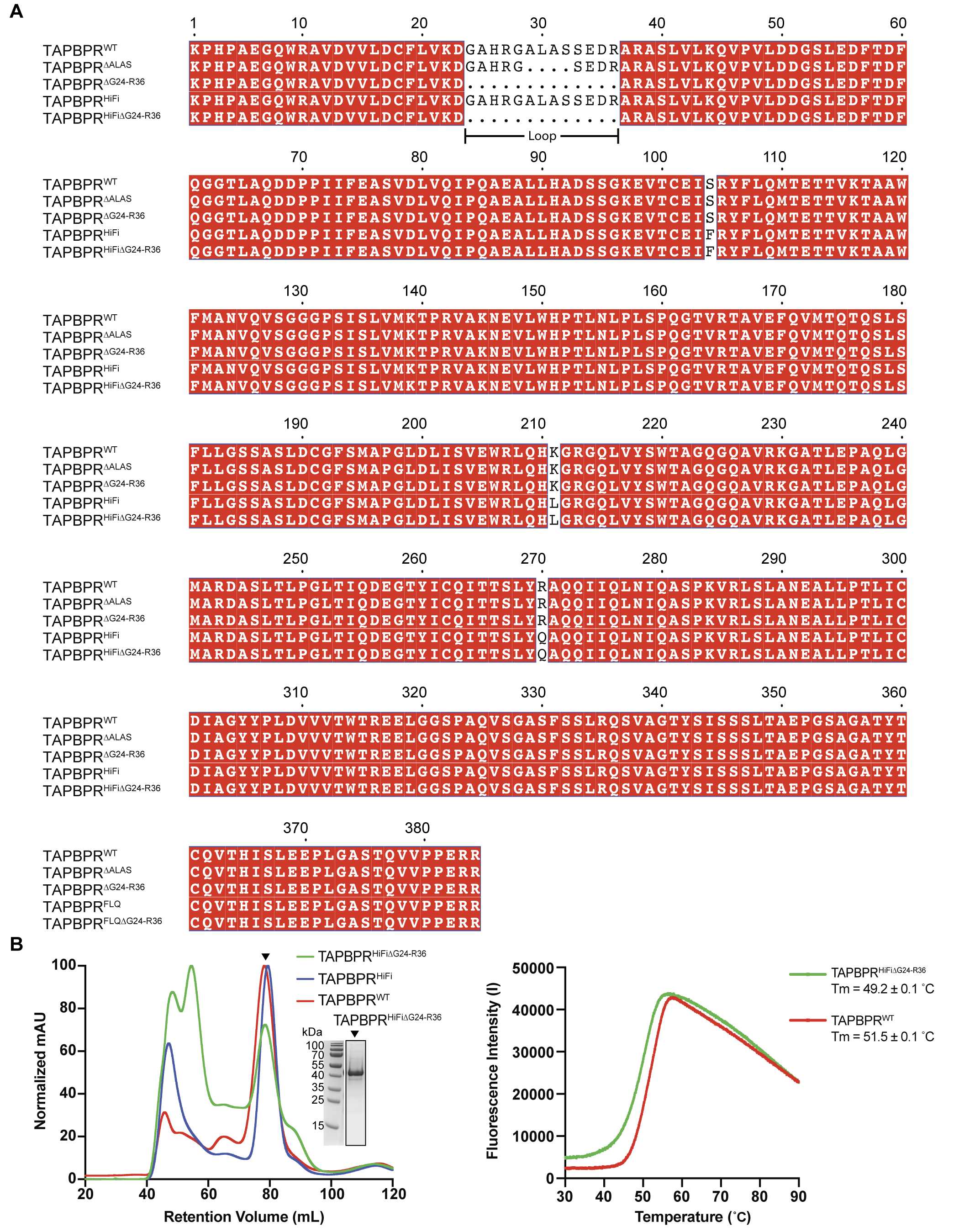
Figure S1. Designed TAPBPR variants used in this study.

(A) Sequence alignment of the human wild-type TAPBPR^WT^, the loop-shorten mutant TAPBPR^∆ALAS^, the loop-deleted mutant TAPBPR^∆G24-R36^, the engineered variant TAPBPR^HiFi^, the loop-deleted variant TAPBPR^HiFi∆G24-R36^. The loop region and amino acid mutations are indicated. Alignments were performed using ClustalOmega^20^ and processed with ESPript 3^21^.

(B) SEC trace of TAPBPR^HiFi∆G24-R36^ mutant (green). The arrow indicates the protein peak relative to TAPBPR^WT^ (red) and TAPBPR^HiFi^ (blue), which is further confirmed by SDS/PAGE analysis. DSF of TAPBPR^HiFi∆G24-R36^ (Tm = 49.2 ˚C, green) relative to TAPBPR^WT^ (Tm = 51.5 ˚C, red).

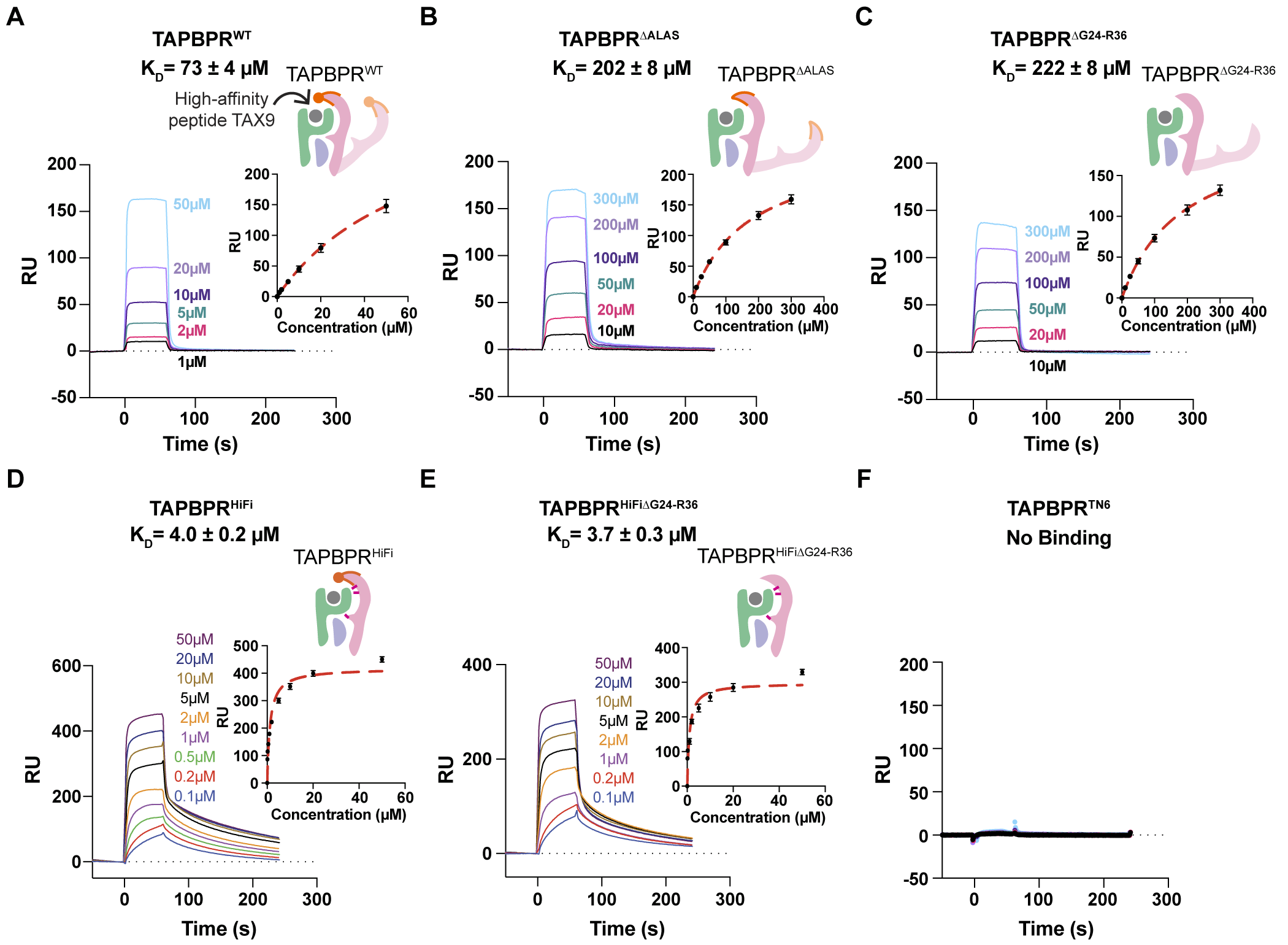
 Figure S2. Surface Plasmon Resonance on direct interactions between TAX9/HLA-A*02:01/β_2_m and different TAPBPR variants.

(A)-(F) Representative SPR sensorgrams of various concentrations of TAPBPR^WT^(A), TAPBPR^∆ALAS^ (B), TAPBPR^∆G24-R36^ (C), TAPBPR^HiFi^ (D), TAPBPR^HiFi∆G24-R36^ (E), and TAPBPR^TN6^ (F) flowed over a streptavidin chip coupled with TAX9/HLA-A*02:01 in molar excess TAX9 peptide. The concentrations of analyte are noted. The equilibrium constant, K_D_, is the mean ± SD for n = 3 independent experiments. The K_D_s of TAPBPR^WT^, TAPBPR^∆ALAS^, and TAPBPR^∆G24-R36^ are estimated values since the saturation of binding was not reached.

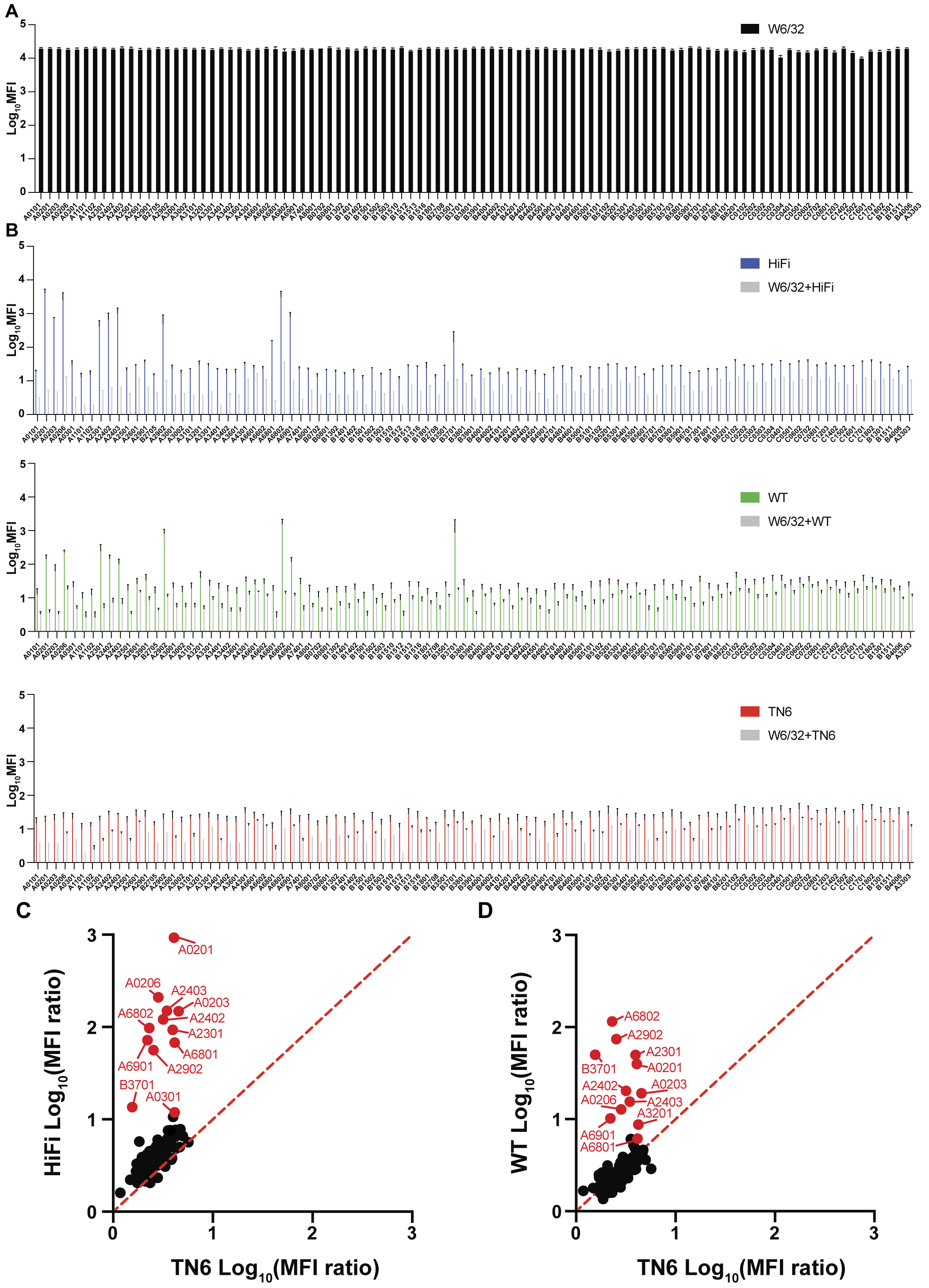

Figure S3. Binding levels of TAPBPR tetramers on HLA single antigen beads.

(A) Levels of folded MHC-I molecules captured on single antigen beads (SABs), related to Fig. 1F, were detected using the primary anti-HLA Class I antibody W6/32 (Abcam, ab22432) and the secondary anti-mouse PE-conjugated antibody (Abcam, ab97024). Similar levels of peptide-loaded MHC-I molecules were observed across different HLA allotypes.

(B) Bar graphs showing the Log_10_ Mean Fluorescence Intensity (MFI) levels of tetramerized TAPBPR^HiFi^, TAPBPR^WT^, and TAPBPR^TN6^ binding to HLA molecules on SABs. Tetramer staining upon the W6/32 antibody incubation, shown in grey, was used to control for background staining levels. The plotted data were generated based on n=2 or 3 independent experiments, and the standard deviation is depicted as error bars.

(C)-(D) Correlation of Log_10_ (MFI ratio) for TAPBPR^HiFi^ (C) and TAPBPR^WT^ (D) relative to TAPBPR^TN6^ plotted in fig. S3B. Dots falling on the dashed red line represent HLA allotypes demonstrating the same MFI levels upon incubation with TAPBPR^WT^ or TAPBPR^HiFi^ tetramers relative to TAPBPR^TN6^.

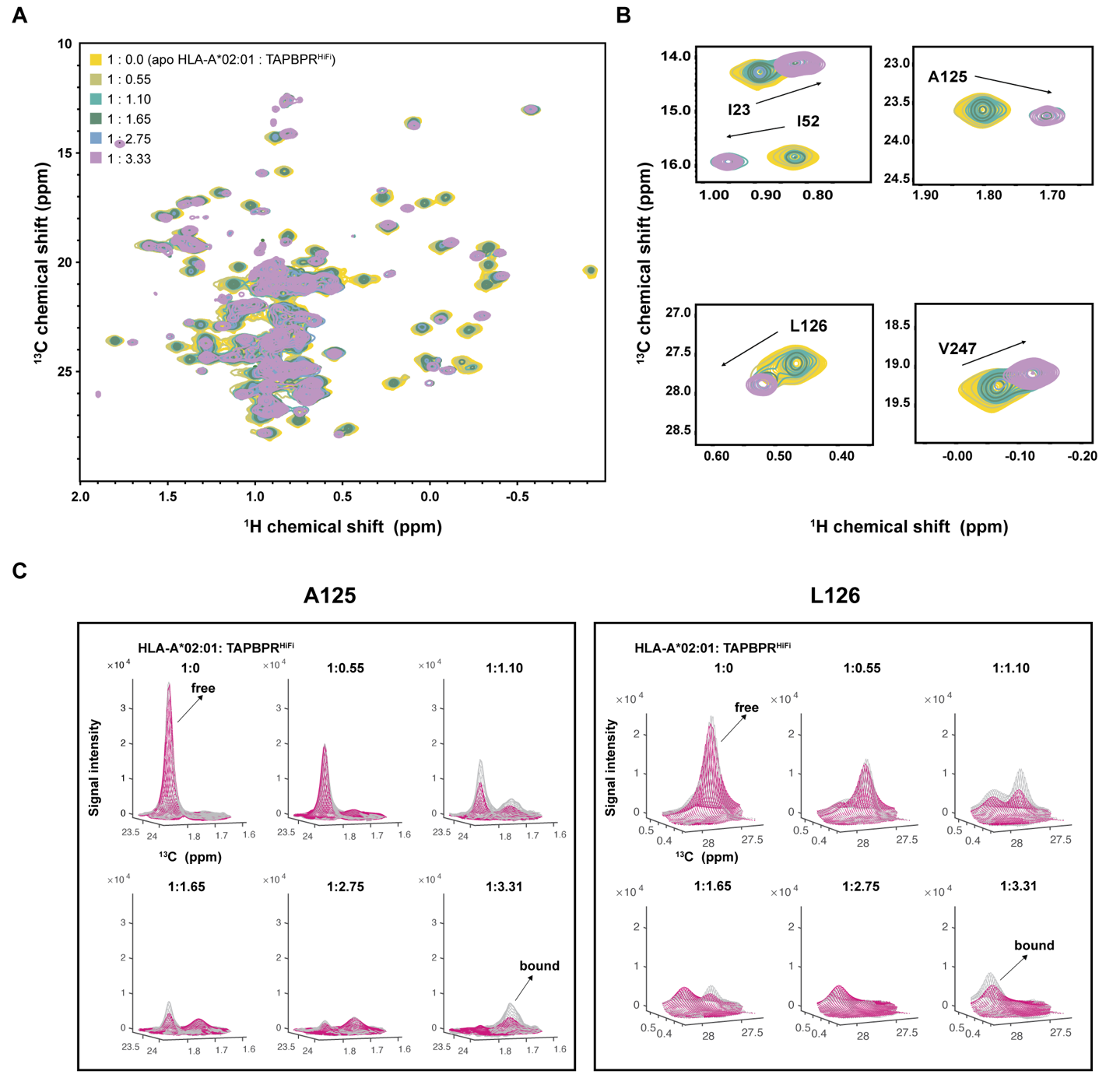
Figure S4. NMR spectra overlay and line shape analysis of peptide-loaded HLA-A*02:01 upon TAPBPR^HiFi^ titration.

(A) 2D ^1^H-^13^C HMQC spectral overlay of selectively ^13^C/^1^H AILV methyl-labeled HLA-A*02:01/TAX9/b_2_m (labeled at the heavy chain) on a ^12^C/^2^H background titrated with TAPBPR^HiFi^ at natural isotopic abundance in the molar ratios: 1: 0 (yellow), 1: 0.55 (khaki), 1: 1.10 (cyan), 1: 1.65 (aquamarine), 1: 2.75 (light blue), 1: 3.31 (purple).

(B) NMR resonances for selected methyl groups (I23, I52, A125, L126, V247) undergoing slow exchange between the free and TAPBPR^HiFi^-bound states.

(C) Representative NMR line shape fitting (shown in magenta) for the resonances of heavy chain methyl from residues A125 and L126 shown on the experimental 2D line shape (grey). All fits were obtained using TITAN^7^.

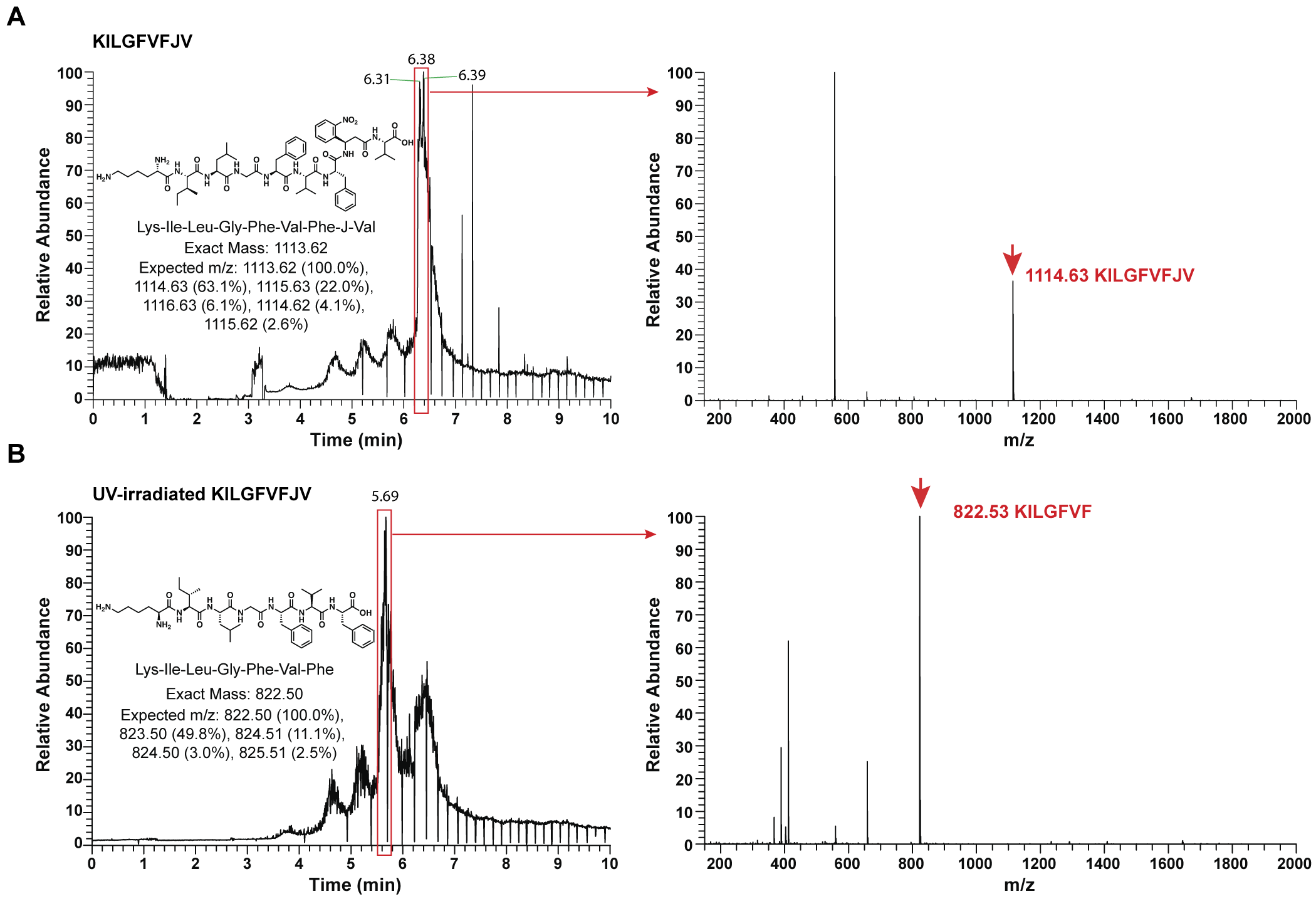

Figure S5. LC-MS validation of UV-irradiated peptide ligand.

(A)-(B) LC-MS analysis of peptide KILGFVFJV (A) and UV-irradiated peptide KILGFVFJV (B). J = 3-amino-3-(2-nitrophenyl)-propionic acid. UV irradiation was performed at 365 nm for 40 minutes at 4˚C. Left panel: the LC chromatogram trace of each sample. Right panel: relative abundance for the selected time interval (red box). In (A), the presence of KILGFVFJV is noted (observed and expected mass-to-charge ratios are 1114.63 and 1114.63 m/z, respectively). In contrast, UV irradiation of KILGFVFJV in (B) results in a peptide fragment KILGFVF (observed and expected mass-to-charge ratios are 822.

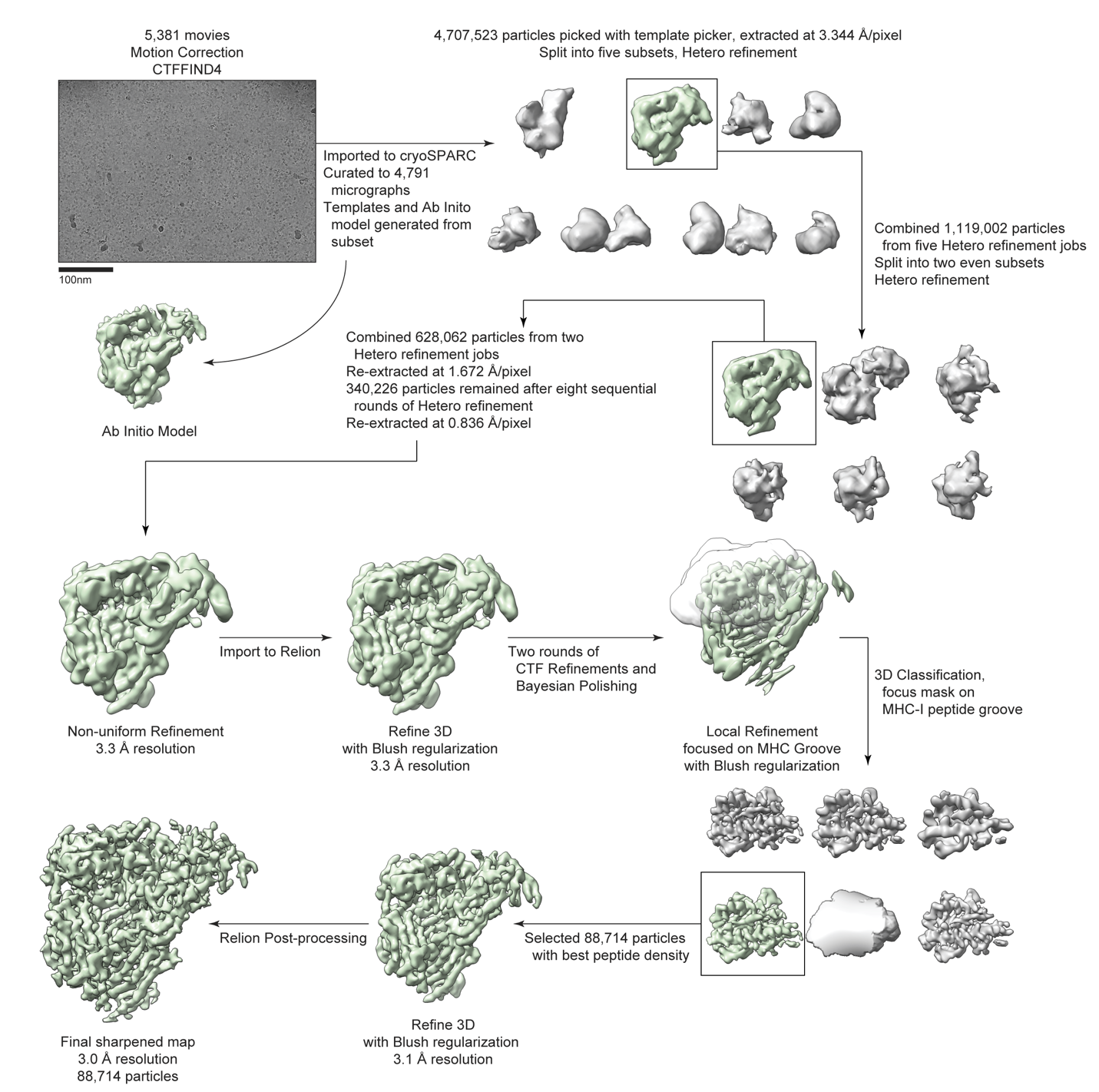
 Figure S6. Cryo-EM data processing workflow.

Cryo-EM data processing workflow including a representative micrograph (top left).

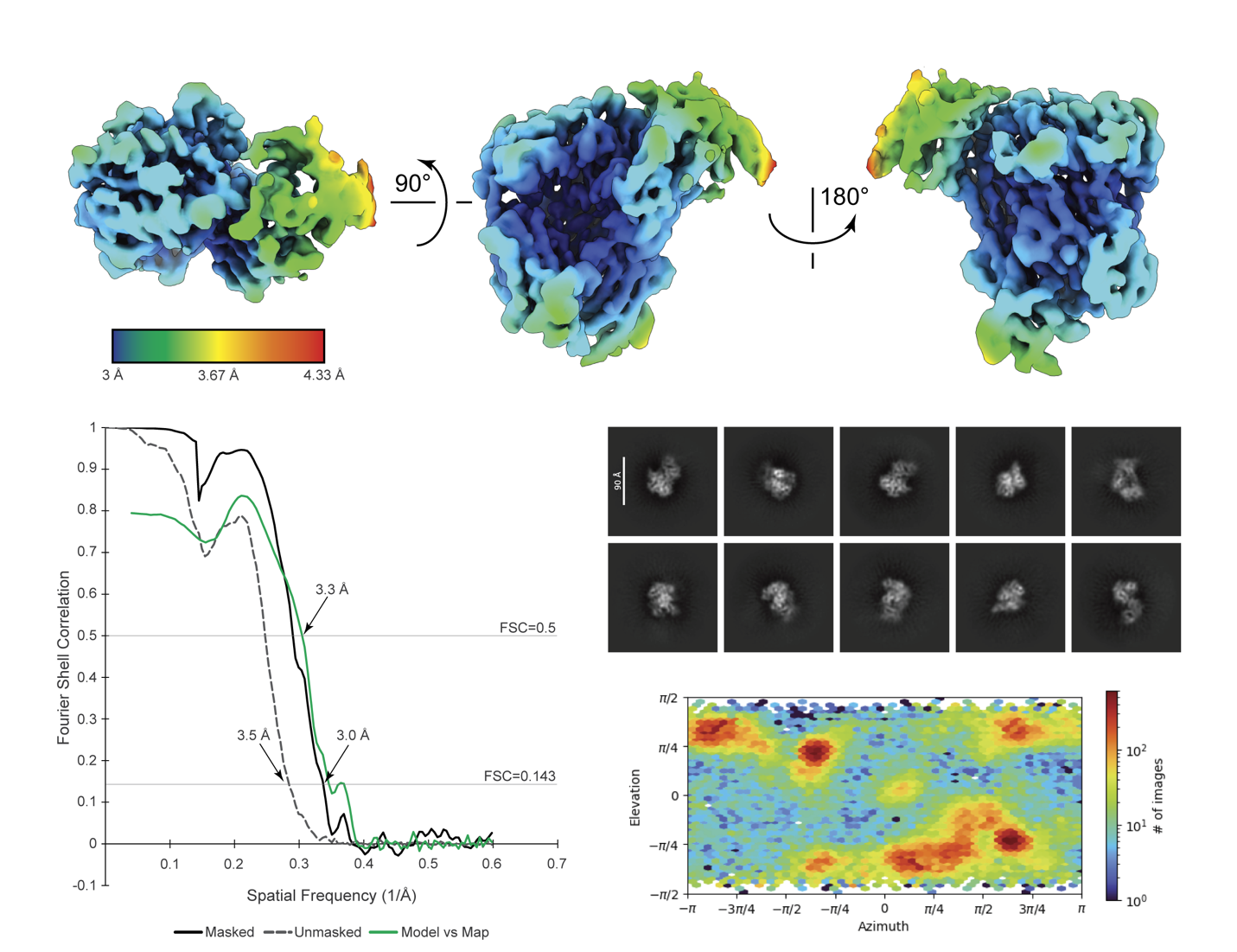

Figure S7. CryoEM data quality.

Local resolution is calculated using Relion’s^14^ local resolution tool (top panels). Unmasked (grey dashed), masked (black), and movel versus map (green) Fourier shell correlation (FSC) curves (bottom left panel). Representative 2D classes generated in cryoSPARC^9^ from final particles (middle right panel) and angular distribution from cryoSPARC reconstruct only based on Relion angular assignment (bottom right panel).

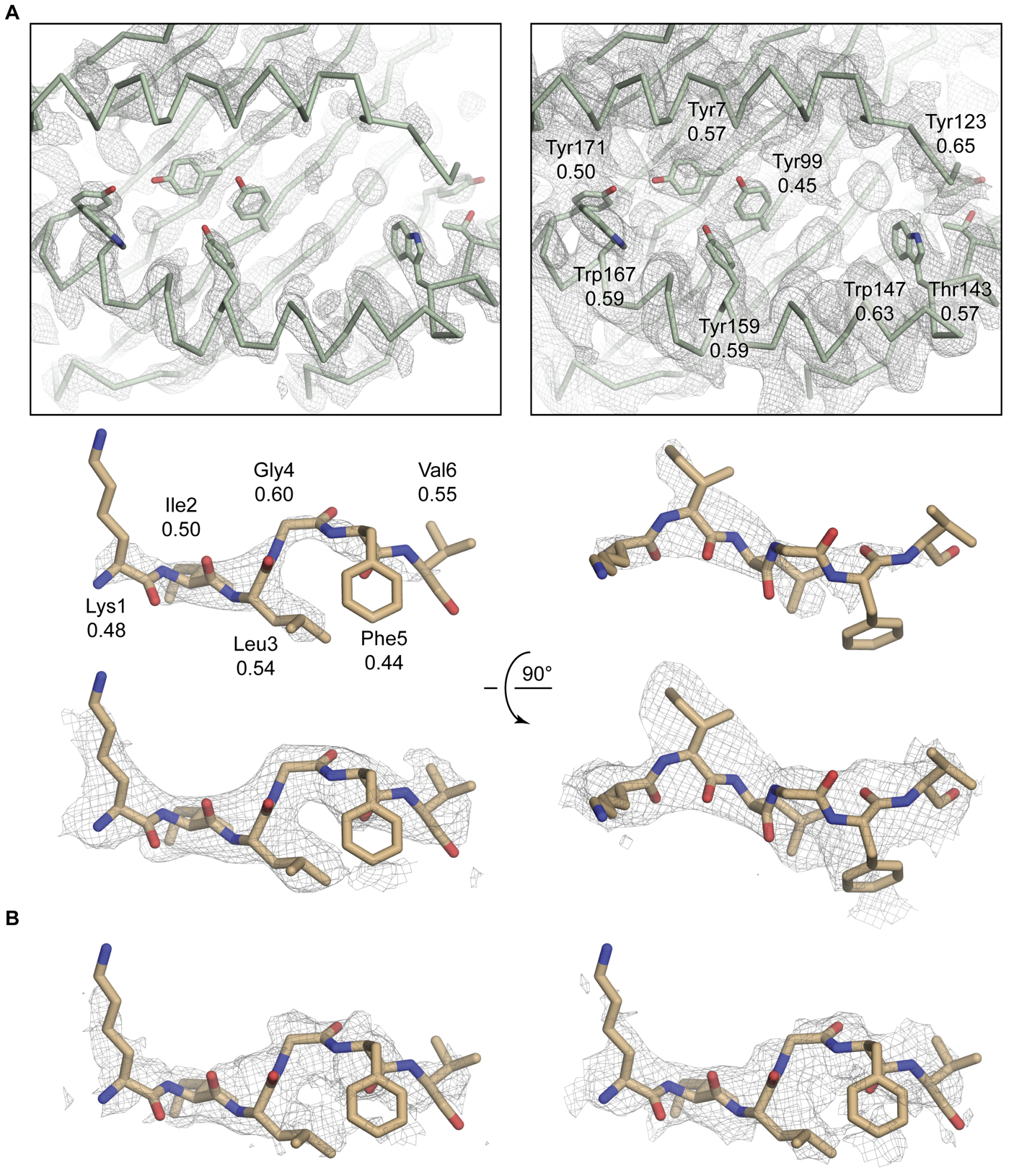
Figure S8. Fit quality of peptide in the MHC-I groove.

(A) Map density and full residue Q-scores for the key residues of the MHC-I groove (top panels) and the peptide (bottom panels). The map is contoured at σ = 10 (top left, middle panels) or σ = 5 (top right, bottom panels). The overall average Q-score for the full structure is 0.52, and the expected average Q-score in a map at 3.0 Å resolution is 0.49.

**(B)** Half map density for the peptide. Peptide density for half map 1 (left) and half map 2 (right). Maps are contoured at σ = 5.

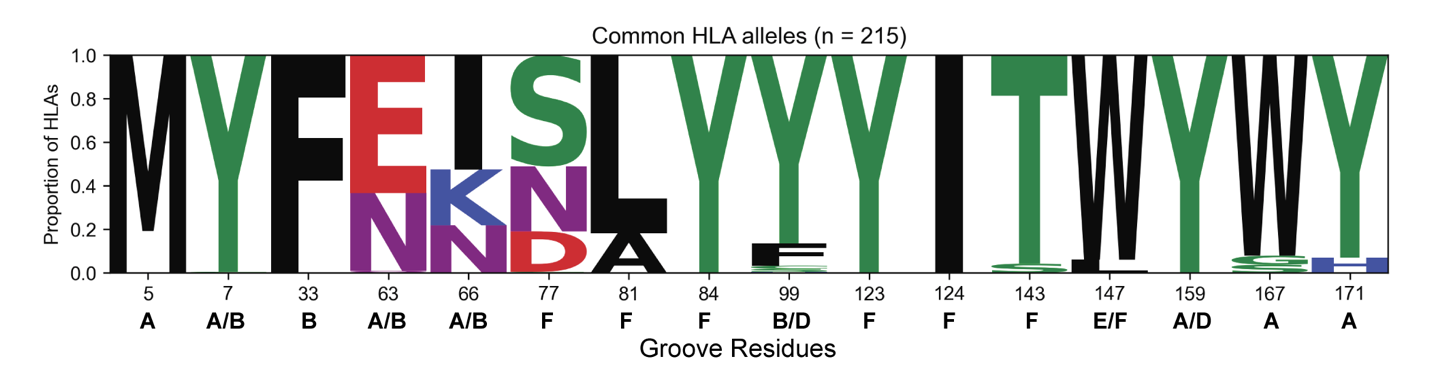
 **Figure S9. Sequence logo of MHC-I heavy chain residues in the A-, B-, D-, E-, and F-peptide-binding pockets.**

Seq2logo visualization^22^ depicting the degree of polymorphism for different MHC-I peptide binding groove residues computed using 215 common HLA-A*, -B*, and -C* allotypes. The location of each residue position in the HLA peptide binding pockets^23^ is noted.

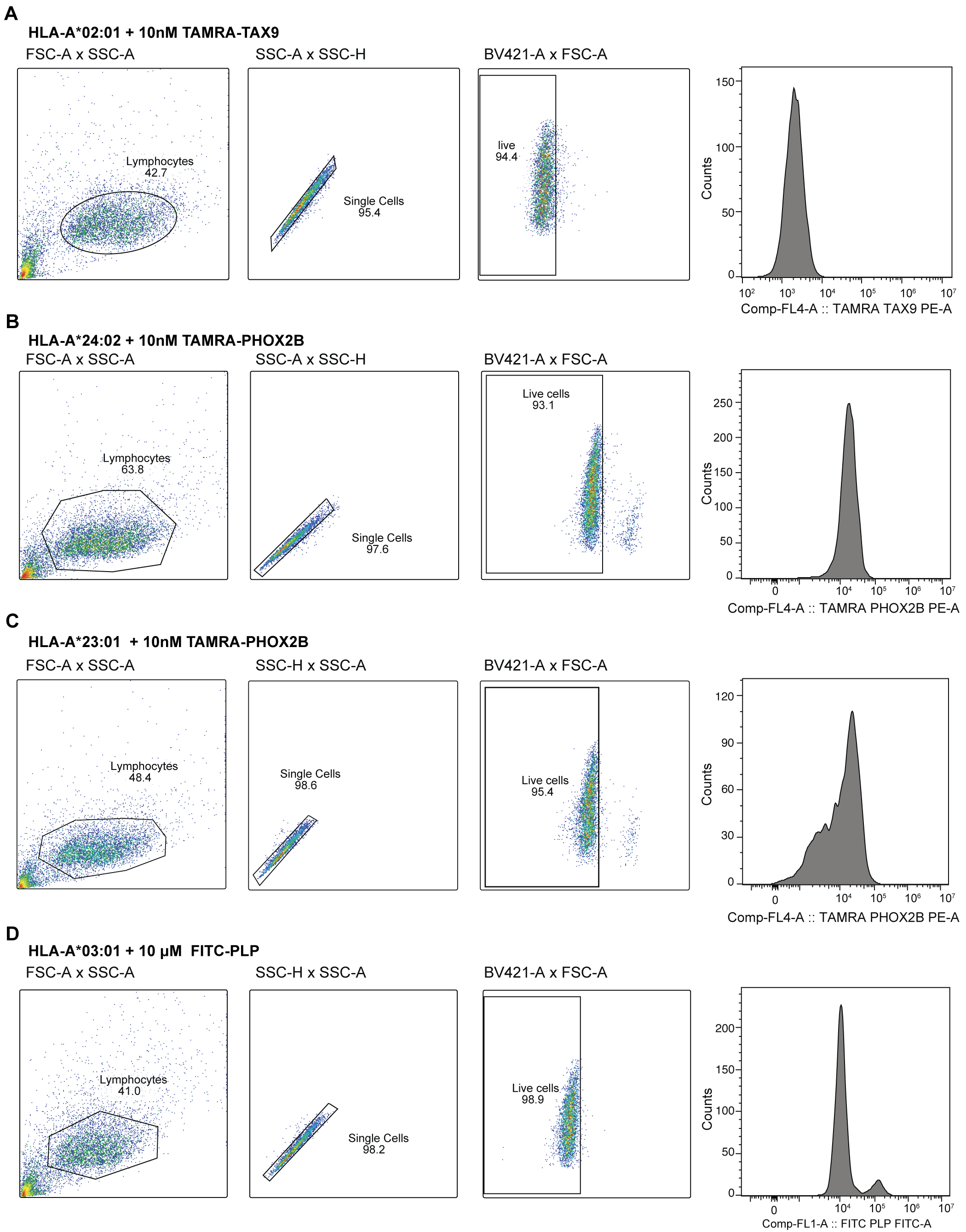
 Figure S10.Representative flow cytometry gating strategy of monoallelic 722.211 cell lines.

(A)-(D) Monoallelic HLA-A*02:01 (A), 24:02 (B), 23:01 (C), or 03:01 (D) 722.211 cell lines were thawed and recovered before performing peptide exchange. Cells were sorted by side and forward scatter (FSC-A and SSC-A) followed by single cell isolation (SSC-A and SSC-H). Gating for live cells was determined by LIVE/DEAD™ Fixable Violet Dead Cell Stain Kit. Gates are shown in the black box, and the percentages of events are gated in parentheses. Cells incubated with no chaperone and corresponding peptide, as indicated, are shown. The acquisition was performed on CytoFLEX LX (Beckman Coulter), and the data were analyzed by FlowJo v10.10.0.

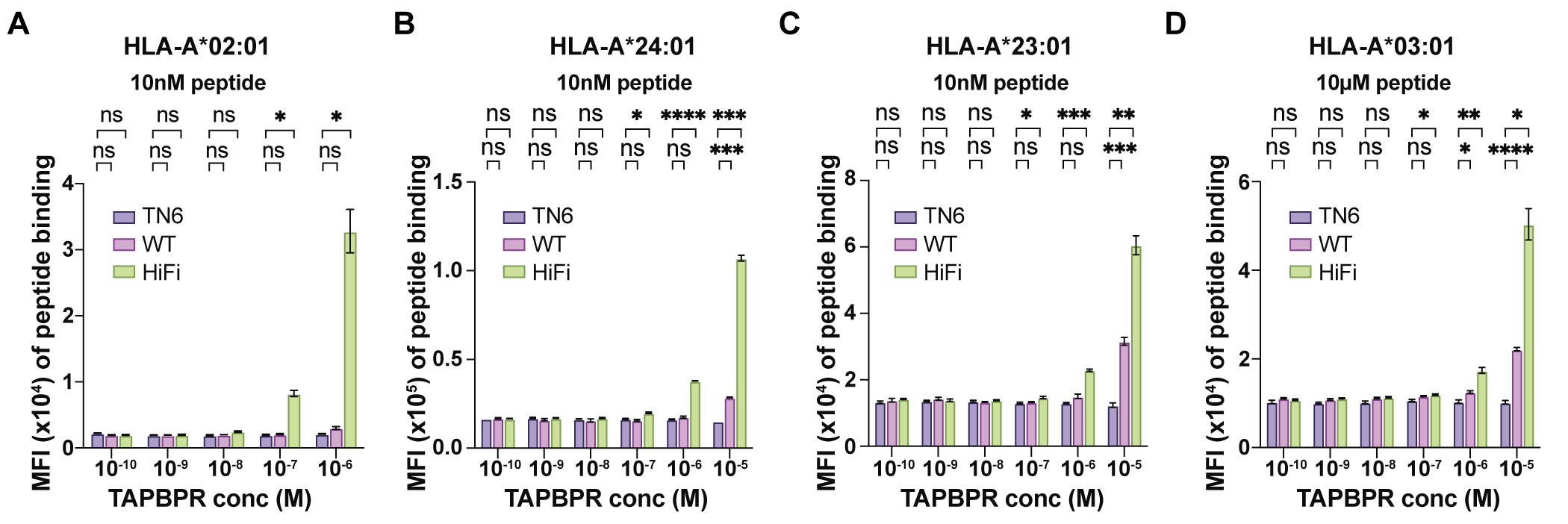
Figure S11. Soluble TAPBPR at different concentrations mediated peptide exchange on surface MHC-I.

(A)-(D) Bar graph summarizing the median fluorescence intensity (MFI) of fluorescent peptide binding to monoallelic HLA-A*02:01 (A), A*24:02 (B), A23:01 (C), and A*03:01 (D) 721.211 cell lines in the presence of soluble TAPBPR at different concentrations, as indicated, from 3 independent experiments. HLA-A*02:01, A*24:02, and A23:01 721.211 cell lines were incubated in the presence of soluble TAPBPR^TN6^, TAPBPR^WT^, and TAPBPR^HiFi^ for 15 minutes at 37 °C, followed by incubation with 10 nM fluorescent peptide for 60 minutes. HLA-A*03:01 monoallelic cell line was incubated in the presence of soluble TAPBPR^TN6^, TAPBPR^WT^, and TAPBPR^HiFi^, followed by incubation with 10 μM fluorescent peptide for 60 minutes. Two-way ANOVA was performed relative to the TAPBPR^TN6^, P > 0.0.1234 (ns), P < 0.0332(*), P < 0.0021(**), P < 0.0002(***), and P < 0.0001(****).

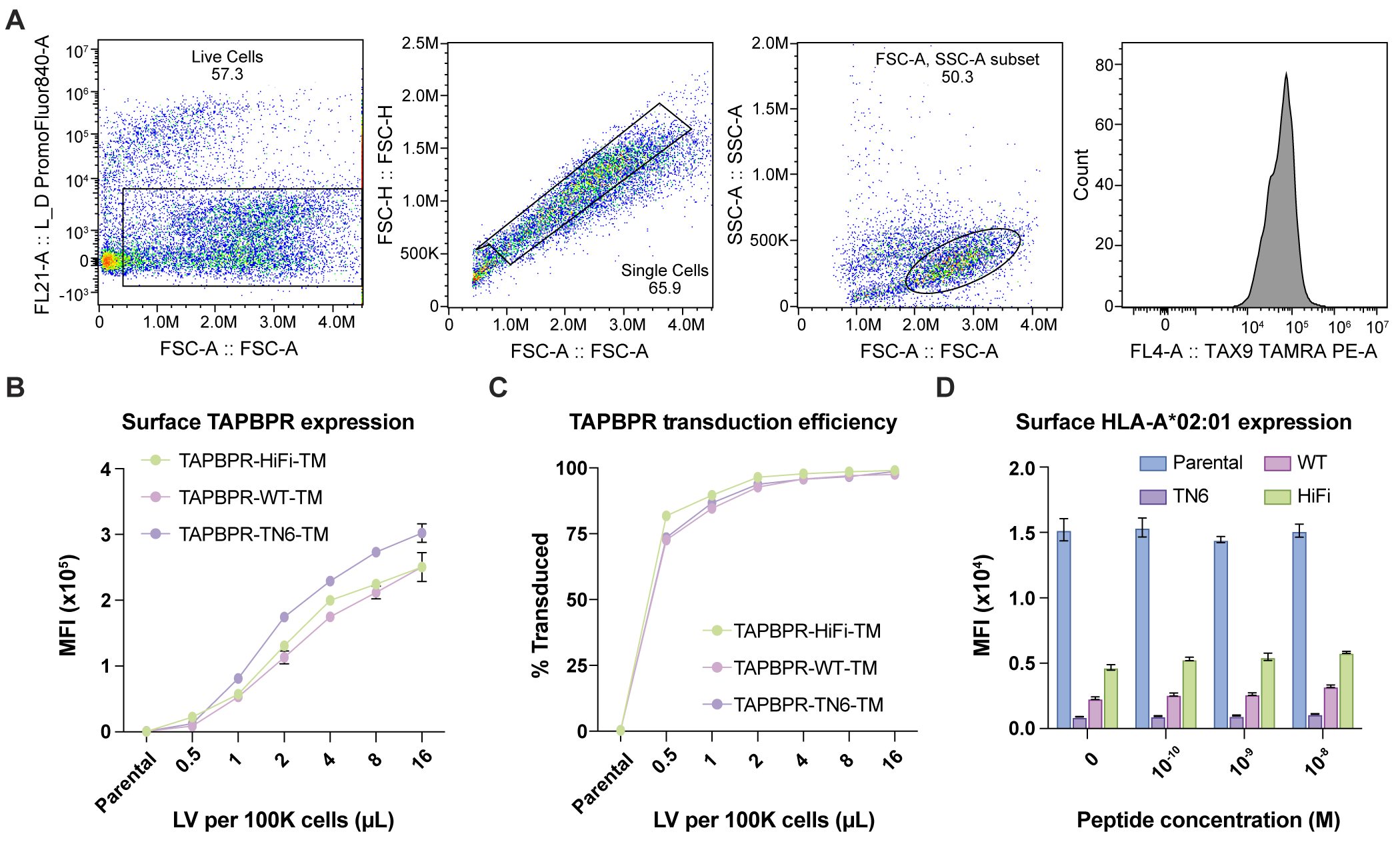
Figure S12. Flow cytometry analysis of T2 cell lines.

(A) Representative flow cytometry gating strategy of T2 cell lines in peptide exchange experiments. Cells were incubated with varying concentrations of fluorescent TAMRA-TAX9 peptide for 60 minutes before washing with FACS buffer, staining with LIVE/DEAD™ Fixable Near IR (876) Viability dye, and fixation with 4% PFA in PBS. Analyzed cells were gated on live singlets of appropriate size. Cells incubated with 10^-5^M of TAX9-TAMARA are shown. The acquisition was performed on CytoFLEX LX (Beckman Coulter), and the data were analyzed by FlowJo v10.10.0.

(B) Surface TAPBPR expression of TAPBPR^HiFi^, TAPBPR^WT^, and TAPBPR^TN6^-TM transduced T2 cell lines, as measured by anti-FLAG MFI.

(C) TAPBPR transduction efficiency of TAPBPR^HiFi^, TAPBPR^WT^, and TAPBPR^TN6^-TM transduced T2 cell lines relative to the non-transduced parental cell line. Cells with similar transduction efficiency and levels of FLAG expression were selected for downstream peptide exchange assays.

(D) Surface HLA-A*02:01 expression of TAPBPR^HiFi^, TAPBPR^WT^, and TAPBPR^TN6^-TM transduced and parental T2 cell lines from cells selected for peptide exchange assays.

Table S1.

**Dissociation equilibrium constants of TAPBPR^HiFi^ binding to TAX9/HLA-A*02:01/β_2_m fitted from NMR line shape analysis of different methyl resonances.** The NMR line shapes of 2D ^13^C/^1^H HMQC resonances corresponding to different methyl-bearing sidechains on HLA-A*02:01 undergoing slow exchange between the free and TAPBPR^HiFi^-bound states were fitted individually to obtain dissociation equilibrium constants, K_D_. The globally fitted value was 13.9 ± 0.7 μM. Line shape analysis was performed using TITAN ^7^ with bootstrap error analysis using 100 replicas.

| Residue | K_D_ (μM) |
| --- | --- |
| I23δ1 | 11.7 ± 0.3 |
| V34γ2 | 17.4 ± 0.9 |
| A125Cβ | 13.8 ± 0.4 |
| V103γ2 | 10.5 ± 0.6 |
| L126δ2 | 14.2 ± 0.5 |
| A117Cβ | 13.9 ± 1.1 |
| L156δ2 | 12.5 ± 0.6 |
| V247γ1 | 16.9 ± 0.8 |
| V247γ2 | 13.1 ± 0.4 |

Table S2.

CryoEM and structural refinement statistics.

|  | pMHC-I/TAPBPR complex  EMD-45360  PDB 9C96 |
| --- | --- |
| **Data Collection and processing** | |
| Magnification | 105,000x |
| Voltage (kV) | 300 |
| Electron exposure (e^-^/Å^2^) | 40.5 |
| Defocus range (μm) | -0.8 to -3.0 |
| Pixel size (Å/pixel) | 0.836 |
| Micrographs (no.) | 5,381 |
| Particles picked (no.) | 4,707,523 |
| Final particles (no.) | 88,714 |
| Map sharpening *B* factor (Å^2^) | -78 |
| Map resolution (masked/unmasked) (Å) | 3.0/3.5 |
| FSC threshold | 0.143 |
| **Model Refinement** | |
| Model resolution cut-off, FSC 0.5 (Å) | 3.3 |
| Model Composition | |
| Non-hydrogen atoms | 4,687 |
| Protein residues | 595 |
| MolProbity score | 1.77 |
| Clashscore | 6.86 |
| Rotamer outliers (%) | 1.59 |
| Cβ outliers (%) | 0.00 |
| CαBLAM outliers (%) | 0.95 |
| R.M.S. deviations | |
| Bond lengths (Å) (# > 4σ) | 0.004 (0) |
| Bond angles (°) (# > 4σ) | 0.659 (0) |
| *B* factors (Å^2^) | |
| min/max/mean | 6.49/113.30/57.11 |
| Ramachandran plot | |
| Favored (%) | 98.00 |
| Allowed (%) | 2.00 |
| Disallowed (%) | 0.00 |

Movie S1.

Overview of the complex structure between TAPBPR^HiFi^ and HLA-A*02:01 bound to a heptameric peptide decoy (KILGFVF). The apo structure of HLA-A*02:01 bound to a high-affinity peptide (GILGFVFTL, PDB ID 2VLL) is overlaid to highlight structural changes in the MHC-I peptide-binding groove.

Movie S2.

Morph between MHC-I structures with different peptide occupancies (empty TAPBPR/MHC-I complex, PDB ID 5WER; TAPBPR/MHC-I bound to a peptide decoy, this work; high-affinity peptide/MHC-I, PDB ID 2VLL) highlighting key residue interactions in the peptide-binding groove. TAPBPR and β_2_m subunits have been removed, for simplicity.

Data S1.

Raw data, sequences, and uncropped gels presented in main and supplementary figures.
